# Extended-Interval Theta Burst Stimulation Enhances Glutamatergic Plasticity and Antidepressant Effects via Reduced GABAergic Recruitment

**DOI:** 10.1101/2024.08.19.608693

**Authors:** Morteza Salimi, Milad Nazari, Anna Linnik, Shruti Joshi, Jonathan Mishler, Ryan Golden, Maxim Bazhenov, Jennifer Rodger, Sahar Jomehpour, Miranda F. Koloski, Jyoti Mishra, Dhakshin S. Ramanathan

## Abstract

Intermittent Theta Burst Stimulation (iTBS) is a patterned stimulation protocol FDA-cleared to treat depression, yet its outcomes are variable and mechanistically unclear. Here, using a combination of calcium imaging, histology, optogenetics and behavior, we show that one parameter of iTBS, the inter-train interval (ITI) between stimulation trains, plays a critical role in modulating GABAergic (and especially parvalbumin) neuronal activity, modulating subsequent neuronal plasticity and antidepressant effects. Shorter ITI stimulation protocols (4-10Ssinter-train intervals) activate GABA neurons, limiting resulting changes in cortical excitability and plasticity compared to extended interval TBS protocols (eTBS, with a 20s ITI). eTBS also drives the largest changes in synaptic / spine plasticity and leads to rapid and durable antidepressant-like effects after only a single stimulation session. Optogenetic activation of GABAergic neurons during eTBS blocks synaptic plasticity and rapid antidepressant effects. Together, these findings reveal a temporal control principle for TBS-induced cortical plasticity and provides a physiology-based strategy to improve TBS efficacy.

**Highlights:**

- Inter-train interval (ITI) controls excitatory-inhibitory balance during theta burst stimulation
- Short ITIs strongly activate PV interneurons limiting longer-term changes in glutamatergic excitatory plasticity
- Extended ITI (20s in particular) reduces inhibitory activity while maintaining sufficient activation of glutamatergic neurons to promote post-stimulation plasticity
- eTBS produces rapid and durable antidepressant-like effects that are blocked by GABAergic co-activation

**Graphical Abstract:** Inter-train interval (ITI) determines the balance between excitation and inhibition during theta burst stimulation (TBS), an FDA-cleared treatment for depression. **(A)** Short ITIs (4s) drive concurrent glutamatergic and GABAergic activation, with inhibitory dominance during stimulation and suppressed long-term modulation of excitability. Extending the ITI to 20s (eTBS) reduces GABAergic recruitment during stimulation, promoting sustained long-term glutamatergic modulation resulting cortical disinhibition, leading to rapid and durable antidepressant-like effects **(B)**. Optogenetic activation of GABAergic interneurons during eTBS abolishes synaptic and antidepressant effects, demonstrating that reduced inhibitory recruitment during eTBS is required for the rapid and durable behavioral effects observed with that protocol.

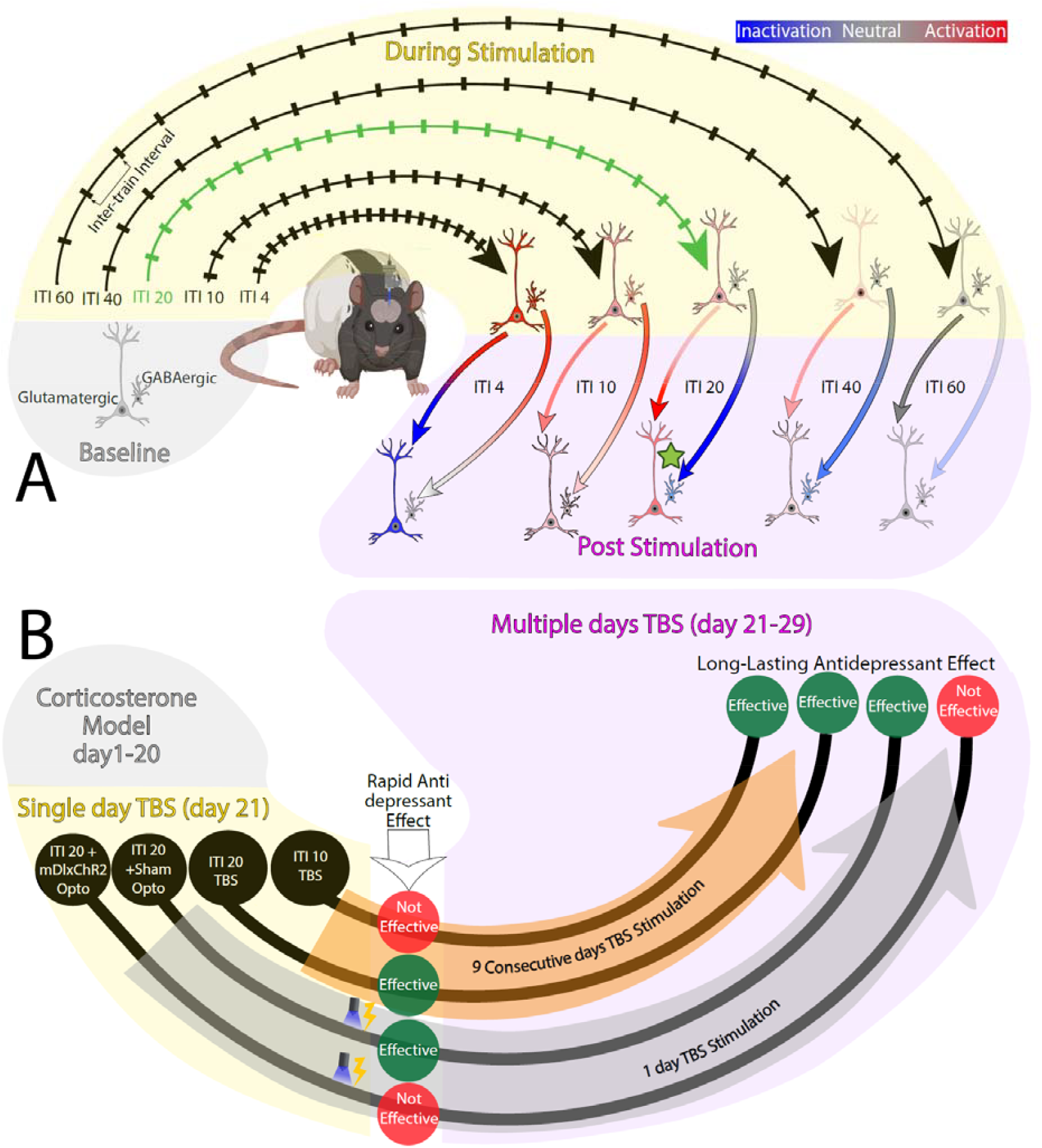

## Introduction

Transcranial magnetic stimulation (TMS) is a noninvasive neuromodulation technique enabling focal cortical activation through magnetic pulses ^1^. Over the past four decades, patterned stimulation protocols have evolved to mimic endogenous brain rhythms and induce lasting plasticity ^2,3^. One of these, theta burst stimulation (TBS), involves brief 50 Hz bursts delivered at a theta rhythm (∼5 Hz). The original TBS protocol, developed and tested in hippocampal slice studies, showed that a single 2s train of electrical TBS stimulation was sufficient to induce long-term potentiation ^4,5^. TBS was adapted for human use, though repeated TBS trains were needed to modulate cortical excitability in vivo ^6^. Importantly, the spacing between trains was shown to be a critical determinant of the direction of excitability changes. Short spacing resulted in a suppression of cortical excitability, while a longer 10s inter-train interval TBS (often called “intermittent TBS”, or iTBS) increased excitability in an LTP-like manner ^6,7^. Subsequent studies confirmed that ITI influences the direction and magnitude of plasticity, though without a clear mechanistic explanation to guide further optimization ^8,9^. Despite this, intermittent TBS has become a cornerstone of translational neurostimulation, both as a tool to probe brain function and as an FDA-cleared treatment for depression ^10,11^.

The circuit-level mechanisms by which intermittent theta burst stimulation shape excitatory-inhibitory dynamics and drives behavioral improvement remain poorly understood. Recent pre-clinical work ^12,13^ showed that plasticity of one particular sub-type of cortical glutamatergic neurons, intratelencephalic neurons, plays a key role in mediating the antidepressant effects of accelerated iTBS. However, these studies did not investigate individual cellular responses to stimulation; the role that GABAergic neurons plan in modulating the effects of TBS, or why differences in ITI during TBS modulates excitability. To address these key questions, we combined in vivo single-cell calcium of glutamatergic and GABAergic neurons during TBS with varying ITIs. We found that shorter ITIs (including both 4 and10s ITI, thus encompassing the standard FDA-cleared iTBS protocol) evoke robust activation in both glutamatergic and GABAergic populations. Shorter ITI protocols with increased GABA recruitment during stimulation show impaired or opposite changes in excitability of glutamatergic neurons post-stimulation. In contrast, an extended interval of 20s between theta burst trains selectively activated glutamatergic neurons with minimal recruitment of fast-spiking GABAergic PV interneurons, resulting in stronger post-stimulation increase in excitability, enhanced circuit- and synaptic-level plasticity and more rapid and durable antidepressant effects. A recurrent neural network model we developed of TBS suggested that enhanced plasticity following extended TBS may be mediated in part by depressed excitatory drive onto inhibitory interneurons, e.g. cortical disinhibition.

These converging cellular, computational and behavioral findings establish the first mechanistic link between stimulation timing, excitation-inhibition balance, and therapeutic efficacy. More broadly, they provide a principled framework for optimizing TBS paradigms to enhance the durability and effectiveness of neuromodulation therapies in neuropsychiatric disorders.

## Results

### Inter-Train Interval Modulates Recruitment and Network Plasticity of Glutamatergic Neurons During Theta Burst Stimulation

To determine how the inter-train interval (ITI) shapes neuronal responses to TBS, we performed in vivo calcium imaging of glutamatergic neurons in the medial prefrontal cortex (Fig 1A, Video S1, Fig S1) using a custom-designed electrolens system described in our recent publication ^14^. Animals received 20 trains of TBS with variable inter-train intervals (ITIs; 4-60 s; Fig 1D and E) using stimulation parameters matched to FDA-cleared clinical iTBS protocols, with the exception that stimulation was delivered electrically rather than magnetically. While we considered utilizing magnetic stimulation for this study, we found it is not feasible with current rodent TMS coils to deliver focal stimulation while also performing imaging with a miniaturized microscope. Moreover, as noted above, the original TBS protocol arose from electrical and not magnetic stimulation ^5^, suggesting that the mode of delivery is less important than the electrophysiological effect on cells. Finally, recent work compared optogenetic and magnetic stimulation ^12,13^ showed surprisingly similar effects, suggesting that the mode of delivery of stimulation is less important than the pattern of stimulation ^15^.

**Figure 1.**
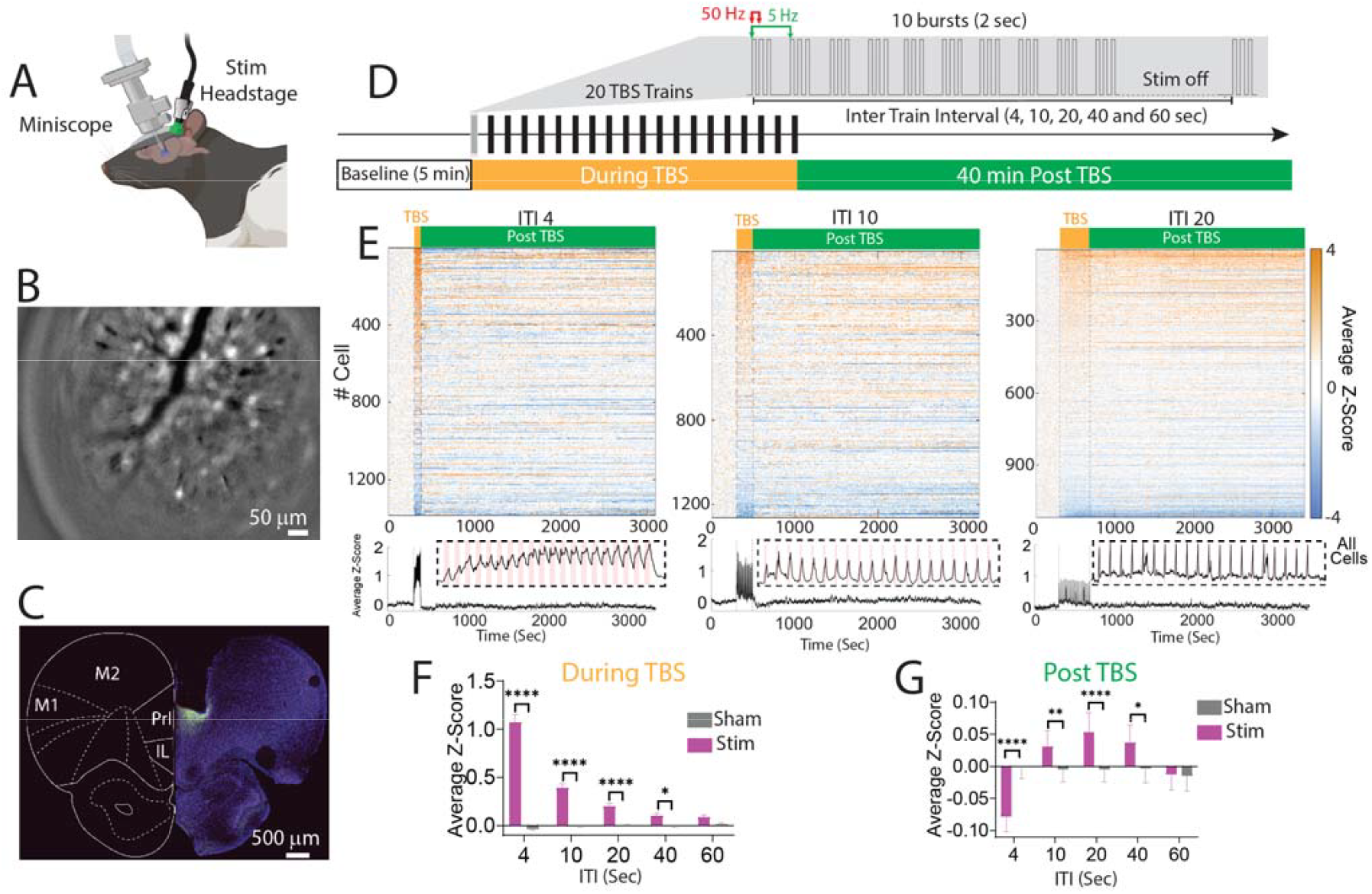
Experimental Design and ITI-Dependent Modulation of All Recorded Glutamatergic Cortical Cell Responses. **(A)** Schematic of the experimental setup. Rats were implanted with micro-wires connected to a headstage adapted for electrical stimulation and an Inscopix GRIN lens for single-cell calcium imaging. **(B)** A representative frame from Inscopix data processing showing spatially mapped cells within the field of view. **(C)** Histological verification of CaMKII-driven GCaMP6 expression in the medial prefrontal cortex (green). **(D)** Theta Burst Stimulation (TBS) was delivered across 20 trains of stimulation. Each train consisted of 30 pulses, with 3 pulses delivered in a 50 Hz burst and bursts repeating at a 5 Hz frequency for 2 seconds. The Inter-train interval (ITI) was varied between sessions (4-60s). Following TBS, calcium imaging continued for 40 minutes. **(E)** Z-scored calcium activity across the entire recording session for all recorded cells individually (top) and averaged across all cells (bottom). Dashed rectangle highl ghts the TBS stimulation window. **(F)**Average calcium activity during stimulation was compared between sham and stimulation conditions across ITIs. Stimulation produced significantly higher calcium activity than sham from ITI 4s through ITI 40s, whereas no significant difference was detected at ITI 60s (see table S1 for more details). **(G)** Post-stimulation activity was analyzed across eight 5-min bins, 40 min total .We found a significantly reduced by stimulation at ITI 4 s, but significantly increased at ITIs 10-40s, with the strongest enhancement observed at ITI 20s; no difference was observed at ITI 60s (see Table S2 and Fig S1 for further details). Data were analyzed two-way ANOVA followed by Sidak’s correction for multiple comparisons. Values are presented as mean ± SEM. *p < 0.05, **p < 0.01, ***p < 0.001, ****p < 0.0001.

To analyze cellular effects while accounting for animal-level variability, we first used a linear mixed-effects model with animal as a random effect (as recommended by ^16^). Stimulation condition, ITI, and their interaction were included as fixed effects and full model with main effects and interactions was performed. The animal-level random effect was not significant (p = 0.179). This model revealed a significant stimulation condition × ITI interaction (F(4, 11879.41) = 83.61, p < 0.001), demonstrating that TBS altered single-neuron calcium activity in an ITI-dependent manner (as this was our primary focus, we did not probe other effects in any significant detail). Group differences between stimulation and sham were followed up within each ITI using a 2-way ANOVA. Stimulation produced significantly higher calcium activity than sham at ITI 4 s (F(1, 11882) = 570.75, p < 0.001), ITI 10 s (F(1, 11882) = 70.22, p < 0.001), ITI 20 s (F(1, 11882) = 16.98, p < 0.001), and ITI 40 s (F(1, 11882) = 5.24, p = 0.022), whereas no significant difference was detected at ITI 60 s (F(1, 11882) = 2.70, p = 0.100; Fig 1F, Table S1). Together, these findings demonstrate that TBS significantly modulates single-neuron calcium responses during stimulation, with the strongest effects occurring at shorter ITIs and diminishing as the interval between trains became longer.

We next asked whether these stimulation-induced changes persist after stimulation. A linear mixed-effects model showed no significant animal level contribution to post-stimulation z-scored calcium activity (p = 0.170). The same model revealed a significant stimulation condition × ITI interaction (F(4, 95148.37) = 16.02, p < 0.001). Using two-way ANOVA to probe post-hoc effects, we found that post-stimulation activity was significantly reduced in the stimulation condition relative to sham at ITI 4s (F(1, 95153) = 35.23, p < 0.001). In contrast, stimulation produced significantly higher post-stimulation activity than sham at ITI 10s (F(1, 95153) = 7.17, p = 0.007), ITI 20s (F(1, 95153) = 17.65, p < 0.001), and ITI 40s (F(1, 95153) = 8.36, p = 0.004). No significant difference was detected at ITI 60s (F(1, 95153) = 0.03, p = 0.860). Modulation followed an upside-down U-shaped curve with the strongest increase observed at ITI 20s (Fig 1G, Table S2).

### Divergent Single Cell Glutamatergic Neuron Responses to Theta Burst Stimulation (TBS)

The core hypothesis underlying TBS is that it induces calcium influx into neurons, triggering intracellular signaling cascades that drive long-term potentiation (LTP) and synaptic strengthening ^4^. Based on this framework, we therefore classified neurons as activated, inactivated, or neutral based on significant deviation from baseline activity during TBS (p < 0.01) and focused our next set of analyses on activated cells (**Fig 2A**). We first quantified the proportion of activated cells during TBS across ITIs (**Fig. 2B** and **C**). Stimulation significantly increased the proportion of activated cells compared with sham at ITI 4s (χ^2^ = 100.03, p < 0.001), ITI 10 s (χ^2^= 42.69, p < 0.001), and ITI 20 s (χ^2^ = 29.22, p < 0.001). No significant difference was observed at ITI 40s (χ^2^ = 0.19, p = 0.661) or ITI 60 s (χ^2^ = 2.87, p = 0.090). Further analyses of inactivated cells are provided in Fig. S2.

**Figure 2.**
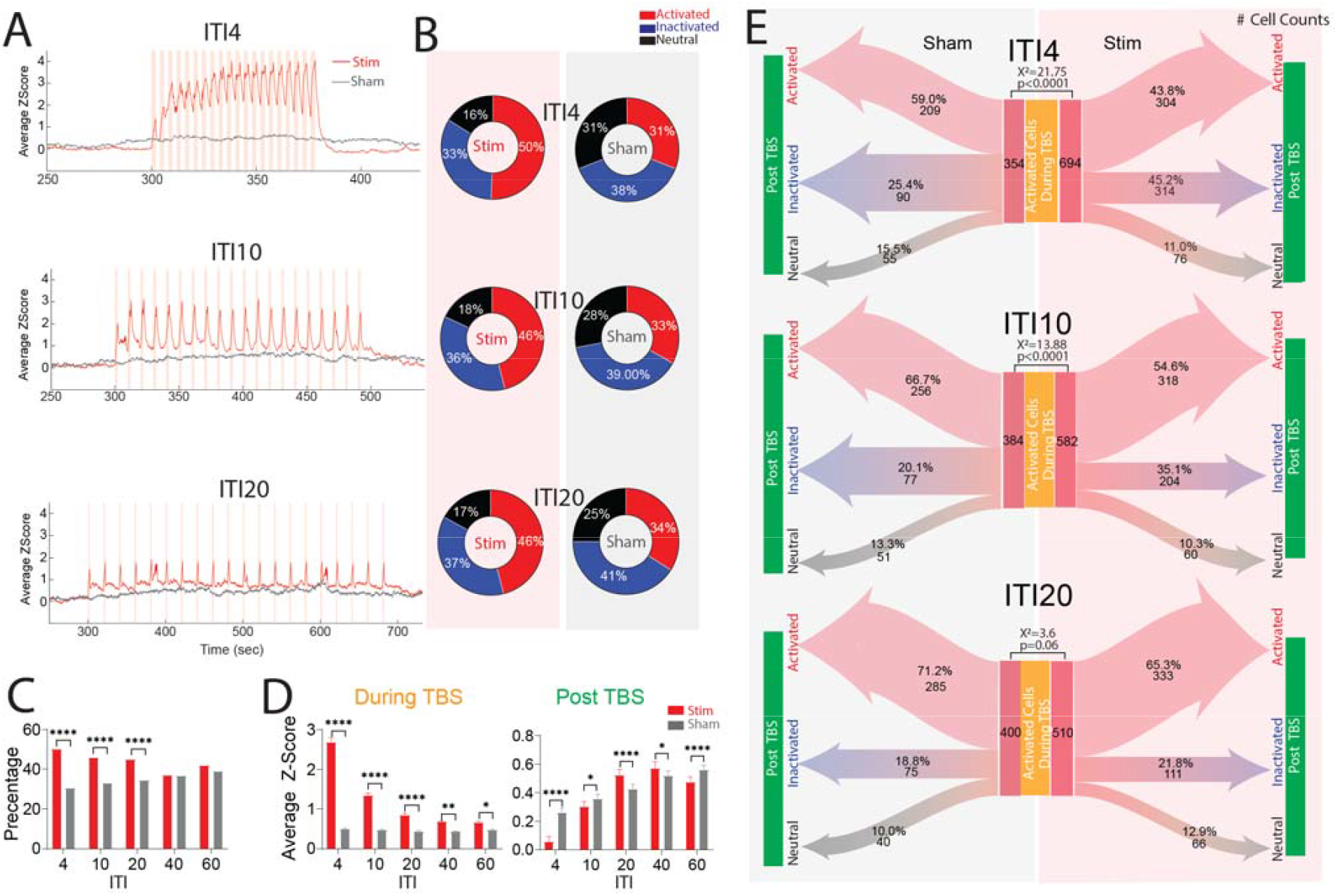
Extended-interval theta burst stimulation stabilizes activated glutamatergic neuronal responses and promotes sustained post-stimulation activity. **(A)** Z-scored calcium activity across the full recording session among cells classified as activated during sham or stimulation conditions. **(B)** Donut plots show the proportion of activated, inactivated, and neutral cells during TBS across ITIs for stimulation and sham conditions. **(C)** The proportion of activated cells during TBS was compared between stimulation and sham conditions across ITIs. Stimulation increased the proportion of activated cells at ITIs 4, 10, and 20s, whereas no difference was observed at ITIs 40 or 60 s. Proportional analyses of inactivated cells are shown in Fig. S2. **(D)** Average calcium activity of activated cells was compared between sham and stimulation conditions during and after TBS. During stimulation, activated cells showed higher calcium activity than sham across all ITIs, with the largest effects at shorter ITIs and progressively smaller effects as ITI increased. Post-TBS activated cells showed lower activity than sham at ITIs 4 and 10s, but higher activity at ITIs 20 and 40 s, indicating that short ITIs induced unstable activation whereas ITI 20s promoted sustained post-stimulation enhancement. **(E)** Transition analysis of activated cells during stimulation was used to compare cells that remained activated post-stimulation with cells that transitioned to either inactivated or neutral states. Short ITIs reduced activated-cell stability, with fewer activated cells remaining activated after stimulation and more cells shifting to inactivated or neutral states at ITIs 4 and 10 s. In contrast, ITI 20s showed a transition pattern more similar to sham, indicating greater preservation of activated-cell stability after stimulation(Fig S4 for ITI 40 and 60s). Data were analyzed using Chi-square tests and two-way ANOVA followed by Sidak’s correction for multiple comparisons. Values are presented as mean ± SEM

We next performed a similar set of analyses in activated cells as we had for the full population of cells in the analysis above; specifically investigating whether there was a significant group × ITI interaction on calcium either during or after TBS (**Fig. 2D**, see **Fig S2** for inactivated cells). During stimulation, a linear mixed-effects model showed no significant animal-level effect (p = 0.169) and a significant stimulation condition × ITI interaction (F(4, 4723) = 94.83, p < 0.001). An analysis of the effects of stimulation within each ITI showed that activated cells had significantly higher calcium activity during stimulation compared with sham at ITI 4s (F(1, 4723) = 634.15, p < 0.001), ITI 10s (F(1, 4723) = 99.17, p < 0.001), ITI 20s (F(1, 4723) = 21.40, p < 0.001), ITI 40s (F(1, 4723) = 7.35, p = 0.007), and ITI 60s (F(1, 4723) = 4.81, p = 0.028), with the magnitude of activation progressively diminishing as ITI increased. Thus, effects during stimulation on activated cells are largely consistent with those observed on the full population. We then asked whether activated cells showed sustained post-stimulation changes. A linear mixed-effects model again showed no significant animal-level contribution for activated cells (p = 0.176) but revealed a significant stimulation condition × ITI interaction (F(4, 37860.24) = 28.23, p < 0.001). Activated cells showed significantly lower post-stimulation activity compared to sham after stimulation at ITI 4s (F(1, 37862) = 82.42, p < 0.001) and ITI 10s (F(1, 37862) = 6.27, p = 0.012), but significantly higher post-stimulation activity at ITI 20s (F(1, 37862) = 16.64, p < 0.001) and ITI 40 s (F(1, 37862) = 4.73, p = 0.030). At ITI 60s, stimulation again produced significantly lower post-stimulation activity compared with sham (F(1, 37862) = 14.22, p < 0.001). See Fig S3 for time bin illustration. This “inverted U” shape was largely consistent with what was observed at the full population level, with peak effects on excitability post-stimulation again observed at the 20S ITI TBS paradigm.

The above analysis suggests that, for shorter duration ITI TBS protocols, the TBS-induced activation is short-lived. To analyze this in a distinct manner, we performed a crosstab analysis to see whether there are differing proportions of activated cells that remain activated post-stim across ITIs (**Fig. 2E**, See **Fig S4** for ITI 40 and 60s). As hypothesized, at both 4s-ITI TBS (χ^2^ = 21.75, p < 0.0001) and 10s ITI TBS(χ^2^ = 13.88, p < 0.0001), there was a reduced proportion of cells that remained activated post-stimulation compared with sham. In contrast, at ITI 20s, the transition pattern was not significantly different from sham (χ^2^ = 3.6, p = 0.06). Together, these results demonstrate that although short ITI TBS paradigms, including 4s and 10s, produce strong activation during stimulation, this activation is unstable and is more likely to shift toward inactivation or neutral states after stimulation. In contrast, though extended intervals show reduced magnitude of acute calcium during TBS, it overall promotes more sustained post-stimulation plasticity.

### Extended Intervals TBS Lessens GABAergic Recruitment

Prior slice studies have shown that excessive GABAergic signaling during TBS trains can block induction of LTP ^17^. As GABAergic neurons, especially fast-spiking parvalbumin (PV) cells, the are effectively when TBS trains are delivered with shorter inter-train intervals ^18^, thus blocking TBS-induced plasticity. To directly test whether shorter ITI TBS protocols enhances GABAergic activity during TBS, we imaged calcium dynamics in mDlx-driven GCaMP-expressing neurons driven by an mDLX promotor, thus increasing expression in putative GABAergic interneurons in the medial prefrontal cortex (**Fig 3A**). Neurons were classified as activated or inactivated using the same criteria applied to glutamatergic cells (**Fig 3B**, Fig S5, Video S2). As hypothesized, short-ITI TBS protocols produced a higher proportion of activated GABAergic cells relative to sham (**Fig. 3B** and **C**). Chi-square analysis showed that stimulation significantly increased the proportion of activated GABAergic cells at ITI 4s (χ^2^ = 96.7, p < 0.0001) and ITI 10s (χ^2^= 14.06, p = 0.0008), but not at ITI 20 s (χ^2^ = 1.09, p = 0.86), ITI 40s (χ^2^ = 6.96, p = 0.053), or ITI 60s (χ^2^ = 1.77, p = 0.69). These findings indicate that closely spaced TBS trains preferentially recruit activated GABAergic interneurons, whereas longer ITIs are less distinct from sham.

**Figure 3.**
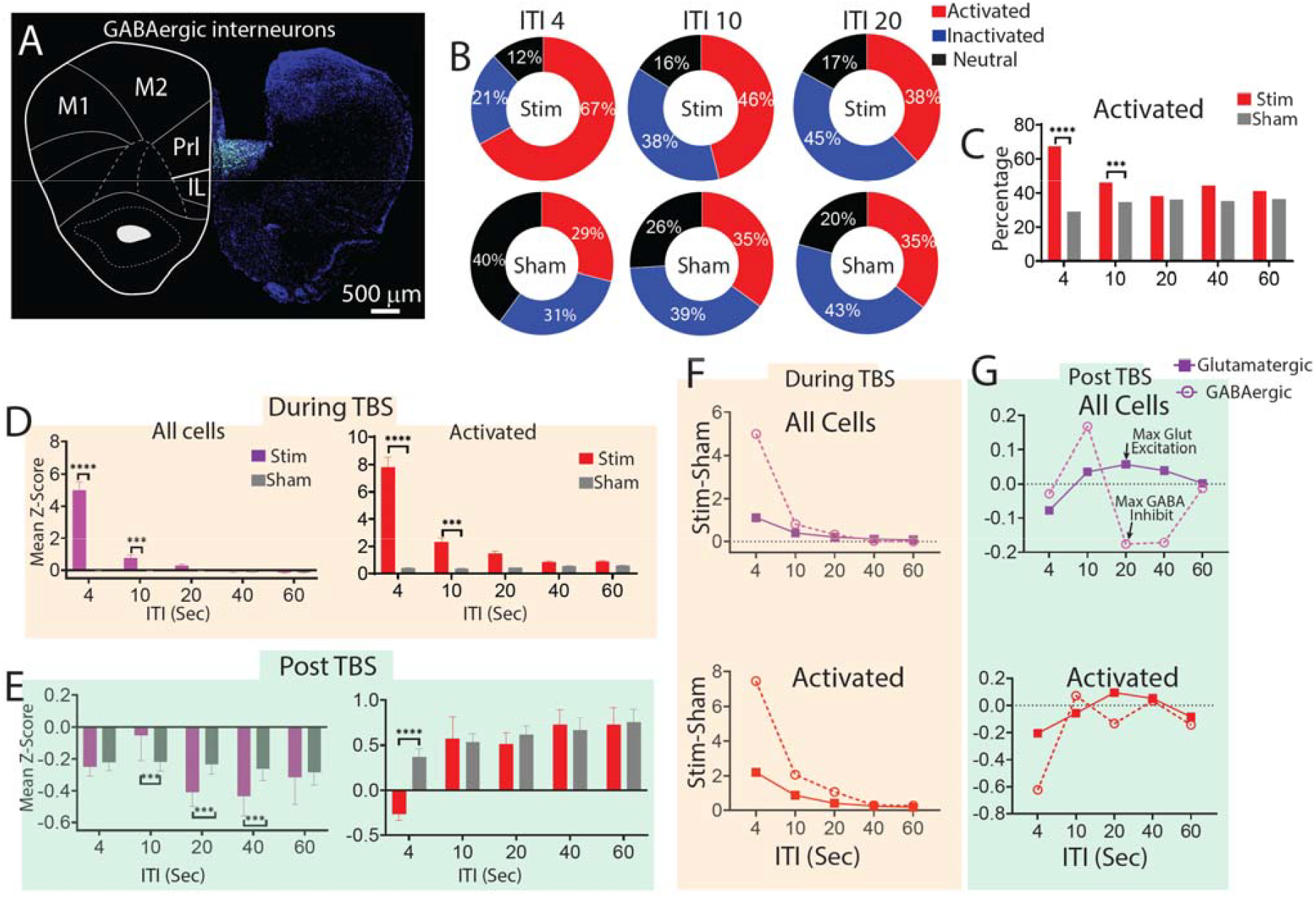
ITI-dependent modulation of GABAergic interneuron activity. (A) Histological verification of mDlx-driven GCaMP expression in medial prefrontal cortex GABAergic interneurons. (B) GABAergic neurons were classified as activated, inactivated, or neutral by comparing mean calcium activity in 4 s time bins around stimulation trains to baseline activity, following the same procedure used for glutamatergic neurons. Donut plots show the proportion of activated, inactivated, and neutral GABAergic cells during TBS across ITIs for stimulation and sham conditions. (C) The proportion of activated GABAergic cells during TBS was compared between stimulation and sham conditions across ITIs. Short ITIs increased the proportion of activated GABAergic cells, with significant increases at ITIs 4 and 10 s, whereas no significant differences were observed at longer ITIs. (D) Average GABAergic calcium activity during stimulation was compared between sham and stimulation conditions across ITIs (Table S3). At the population level, stimulation produced higher calcium activity than sham at ITIs 4 and 10 s, but not at longer ITIs. Among activated GABAergic cells, stimulation similarly increased calcium activity at ITIs 4 and 10 s, with responses diminishing at longer intervals. (E) Post-stimulation GABAergic activity was calculated across eight 5-min bins over the 40-min post-TBS period (Fig S6, Table S4). At the population level, post-stimulation activity was lower than sham at intermediate ITIs, including ITIs 10, 20, and 40 s, but not at ITIs 4 or 60 s. Among activated GABAergic cells, post-stimulation activity was lower than sham only at ITI 4 s, with no significant differences at longer ITIs. (F) Differential plots showing stimulation-minus-sham changes in z-scored calcium activity during TBS. Short ITIs produced the strongest GABAergic recruitment during stimulation, which declined as ITI increased. (G) Differential plots showing stimulation-minus-sham changes in post-stimulation activity. ITI 20 s was associated with maximal glutamatergic excit tion balance. Data were analyzed using Chi-square tests and two-way ANOVA followed by Sidak’s correction for multiple comparisons. Values are presented as mean ± SEM.

We next examined GABAergic calcium activity during stimulation. Linear mixed-effects model showed no significant animal-level effect (p = 0.242) and a significant stimulation condition × ITI interaction (F(4, 3232.12) = 52.22, p < 0.001). Using two-way ANOVA, post hoc comparisons showed that population-level GABAergic activity was significantly higher than sham at ITI 4s (F(1, 3233) = 302.65, p < 0.001) and ITI 10s (F(1, 3233) = 7.66, p = 0.006), but not at ITIs 20–60s. These findings again suggest that short ITIs preferentially recruit GABAergic activity during stimulation, but this effect no longer remains at 20s ITI and above. We also examined just the activated GABAergic cells during stimulation (**Fig. 3D**, Table S3). We found no effect of animal (p = 0.267) and a significant stimulation condition × ITI interaction (F(4, 1323.91) = 19.88, p < 0.001). Post hoc ANOVA showed that stimulation significantly increased calcium activity in activated GABAergic cells at ITI 4s (F(1, 1325) = 132.86, p < 0.001) and ITI 10 s (F(1, 1325) = 8.93, p = 0.003), but not at ITI 20s, 40s, or 60s.

We finally analyzed post-stimulation GABAergic activity, starting at the whole population (**Fig. 3E**; **Fig**. S6; Table S4). We found no animal-level effect (p = 0.179) and a significant stimulation condition × ITI interaction (F(4, 25937.79) = 5.56, p < 0.001). Post-hoc two-way ANOVA demonstrated that post-stimulation GABAergic activity was significantly larger than sham at ITI 10s (F(1, 25941) = 7.20, p = 0.007) but suppressed at ITI 20 s (F(1, 25941) = 7.91, p = 0.005), and ITI 40 s (F(1, 25941) = 7.65, p = 0.006). ITI 4s and ITI 60s did not differ from sham. Next, we examined post-stimulation activity in the activated sub-population of GABAergic cells (**Fig. 3E**). A mixed-effects model showed no significant animal effect GABAergic cells (p = 0.171) and a significant stimulation condition × ITI interaction (F(4, 10667.71) = 6.08, p < 0.001). Using two-way ANOVA, pairwise comparisons showed that activated GABAergic cells had significantly lower post-stimulation activity than sham only at ITI 4s (F(1, 10669) = 37.65, p < 0.001), but no significant differences at ITIs 10–60s. This lack of effect in activated neurons suggests that the effects observed in the full population of GABAergic cells was not a direct effect of TBS but instead an indirect effect mediated by a micro-circuit effect, an idea we explored subsequently with computational modeling.

First, to illustrate the total of these changes across cell-types, we plotted the change in mean glutamatergic and GABAergic activity relative to sham across ITIs, both during and after stimulation (**Fig. 3F–G**). During stimulation, 4s and 10s preferentially recruited GABAergic activity, whereas this preferential modulation recruitment diminished at longer interval TBS. Post-stimulation, ITI-20 and ITI-40 TBS were both associated with glutamatergic excitation and suppression of GABAergic activity. ITI-10, alone, was associated with an overall net positive modulation across both cell-types. Together, these results suggest that short ITIs drive concurrent recruitment of excitatory and inhibitory populations during stimulation, resulting in transient and unstable network responses, whereas extended ITIs reduce inhibitory recruitment during stimulation and support a more favorable post-stimulation excitation/inhibition balance.

### Parvalbumin GABAergic Neurons Are Preferentially Activated by Short ITI TBS Protocols

The physiological data above are limited by the incomplete specificity of the mDLX promoter for GABAergic neurons ^19^ and the lack of resolution across inhibitory neuronal subtypes. To address these limitations, we performed double immunostaining for c-Fos and GAD65 in the prelimbic mPFC (**Fig 4**). Quantification of c-Fos–positive cells alone revealed that short ITIs (4s and 10s) induced a significant increase in neuronal c-Fos activation relative to sham, whereas extended-interval TBS (20s) did not differ from sham across all cells (p=0.146; **Fig 4A-D)**. We quantified GAD-65 expression within c-Fos positive neurons to assess inhibitory activity among the activated cell population. Short ITIs (4s and 10s) produced markedly higher GAD-65 intensity compared to sham and eTBS (20s (ITI 4 s: p < 0.0001 versus sham, p = 0.0003 versus ITI 20 s; ITI 10 s: p = 0.0002 versus sham, p = 0.0059 versus ITI 20s),whereas extended-interval TBS did not significantly differ from sham in GAD-65 intensity (p=0.51; **Fig 4E**), consistent with our in vivo physiology data. We next examined activity within two major interneuron subtypes: parvalbumin (PV)-expressing interneurons (fast-spiking) and somatostatin (SST)-expressing interneurons (slow-spiking) ^18,20^. We quantified intracellular intensity of GAD-65 within individually reconstructed PV and SST cells across ITIs (**Fig 4F**). Two-way ANOVA (stimulation condition × cell type, **Fig 4G**) indicated significant main effects of stimulation condition (F_(1, 16)_ = 128.3, p < 0.0001) and cell type (F_(3, 16)_ = 23.31, p<0.0001) along with a significant interaction (F_(3, 16)_ = 9.497, p=0.0008). Post hoc analysis showed short ITIs robustly increased GAD-65 levels in both cell types, but the effect was significantly greater in PV interneurons, consistent with prior evidence identifying PV cells as primary mediators of fast inhibitory feedback and cortical gain control (Tremblay et al., 2016; Urban-Ciecko and Barth, 2016). Extended-interval stimulation produced no difference from sham in either population, reflecting a restoration of baseline inhibitory tone. To evaluate whether GABA synthesis differed between interneuron subtypes as a function of stimulation interval, we quantified the difference in GAD-65 intensity between PV and SST interneurons (**Fig 4H**). Our one-way ANOVA analysis revealed a significant effect across stimulation conditions (F_(3,16)_ = 9.497, p = 0.0008). Post hoc comparisons showed that short ITIs (4s and 10s) produced markedly greater GAD-65 upregulation in PV relative to SST interneurons compared to sham, indicating enhanced GABA production in PV cells with brief interval TBS. In contrast, extending the ITI to 20s reduced this difference, resulting in values not significantly different from sham, consistent with restoration of baseline inhibitory balance.

**Figure 4.**
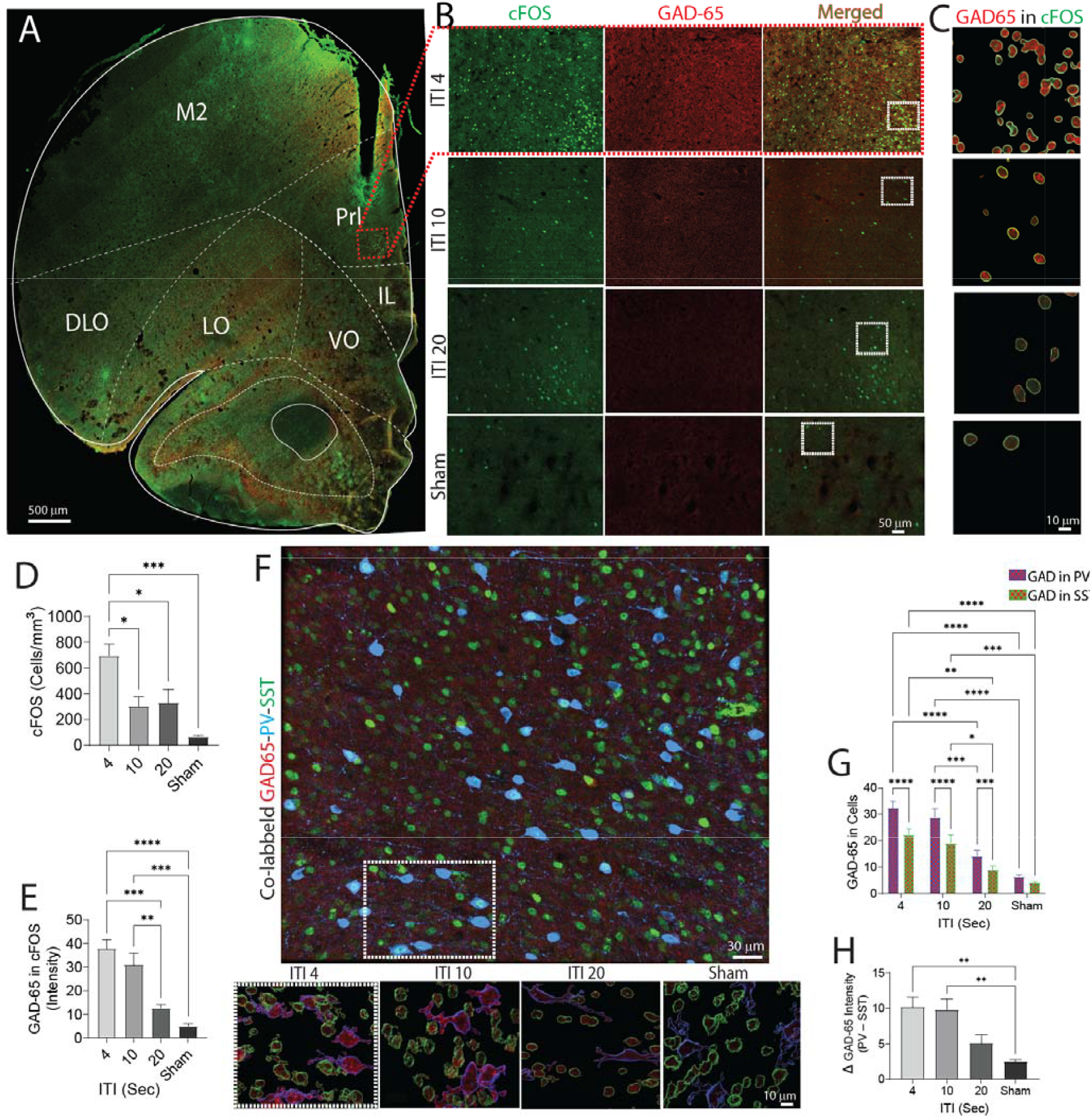
Immunohistochemical analysis of c-Fos and GAD-65 expression in response to different ITIs of theta burst stimulation. **(A)** Representative full-brain section showing double immunostaining for c-Fos (green) and GAD-65 (red) in the prelimbic medial prefrontal cortex (mPFC). All brains were collected 60 min post-stimulation. Sham animals underwent electrode implantation but received no stimulation. **(B)** Magnified view of the prelimbic region from the same section. **(C)** 3D cellular reconstruction generated using Imaris software illustrating GAD-65 (red) expression within c-Fos–positive neurons. **(D)** Quantification of c-Fos-positive cell counts revealed significantly greater activation at short ITIs (4 s and 10 s) compared with sham, whereas extended-interval TBS (20 s) did not differ from sham, indicating reduced neuronal recruitment with longer intervals. **(E)** Mean GAD-65 intensity within c-Fos–positive cells differed significantly across ITIs, with short ITIs (4 s, 10 s) showing markedly higher GAD-65 levels relative to sham and eTBS (20 s). (F) Representative 3D reconstructions showing co-labeling of GAD-65 (red) with parvalbumin (PV, blue) and somatostatin (SST, green) interneurons; lower panels depict examples of GAD-65 localization within individual PV and SST cells. (G) Quantification of GAD-65 intensity within PV and whereas extended-interval TBS (20 s) did not differ from sham. **(H)** Difference in GAD-65 intensity between PV and SST interneurons across stimulation conditions. Short ITIs (4 s and 10 s) showed greater GAD-65 upregulation in PV relative to SST interneurons compared to sham, whereas extending the ITI to 20 s reduced this difference, resulting in values not significantly different from sham. Data are expressed as mean ± SEM (n = 4 rats per group) and analyzed by ANOVA with Sidak’s correction for multiple comparisons. *p < 0.05, **p < 0.01, ***p < 0.001, ****p < 0.0001.

Together, the combined physiologic and histologic reports robustly demonstrate that 20s ITI represents a “sweet spot” with sufficient drive to activate glutamatergic neurons with minimal activation of GABAergic (and particularly parvalbumin) neurons, thus maximally driving cortical in vivo plasticity.

### Spiking Neural Network Model Reveals Disinhibitory Plasticity After Extended-Interval TBS

Our physiological and histological findings suggested that extended-interval TBS (eTBS) induces sustained excitability in glutamatergic neurons and suppression of GABAergic neurons. While the effect on glutamatergic neurons seems directly related to calcium influx (“e.g. activated cells”), the suppression of GABAergic neurons post-stimulation does not occur in cells that were stimulated, suggesting an indirect mechanism. To directly test this, we implemented a spiking recurrent neural network (sRNN) based modeling framework to estimate changes in synaptic weights across neuronal populations following TBS. The sRNN model, implemented with a kernel that converts the spiking activity to calcium fluorescence (see Methods) was trained via backpropagation through time (BPTT) to reconstruct the fluorescence traces of the glutamatergic and GABAergic neurons observed in vivo. The model was first trained on the baseline data (5 min pre stimulation) by optimizing membrane time constants, synaptic weights and synaptic time constants. The model was then further trained on the post TBS data but only the glutamatergic to glutamatergic and glutamatergic to GABAergic synapses were allowed to change and all other parameters were frozen to better isolate our hypotheses of interest. The model successfully reconstructed the in vivo fluorescence for the baseline and post-TBS data and captured the changes in activity of the neurons post TBS compared to baseline (Fig S7).

We next examined the distribution of synaptic weight changes (Δw; 20 min post-stimulation minus baseline) for glutamatergic-to-glutamatergic and glutamatergic-to-GABAergic connections across inter-train intervals (ITIs) (**Fig 5A-B**). Intermediate ITIs (10s and 20s) showed larger changes in connectivity compared to the other ITIs suggesting more circuit restructuring occurred for the intermediate ITIs following TBS. While both connection types exhibited ITI-dependent shifts in synaptic strength, glutamatergic-to-GABAergic connections showed a broader and more negatively shifted distribution, particularly at intermediate ITIs. For our main analysis synaptic weights for GABAergic-to-glutamatergic connections were frozen due to data on inhibitory synaptic plasticity within cortical circuits (however, we found largely similar results when allowing GABAergic to glutamatergic to change).

**Figure 5.**
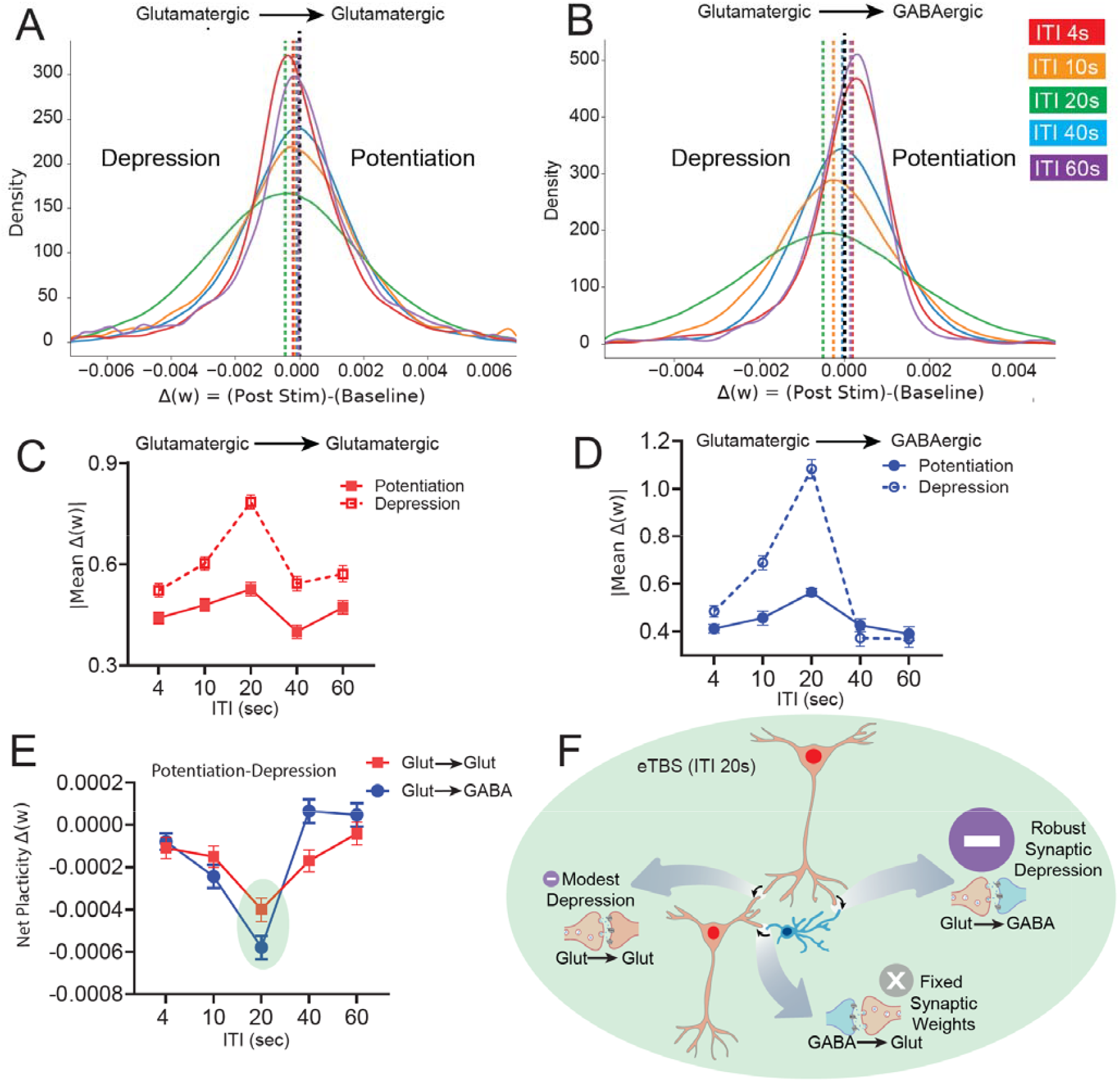
Extended-interval TBS preferentially induces synaptic depression in glutamatergic-to-GABAergic connections. **(A–B)** Distribution of model-derived synaptic weight changes (Δw; 20 min post-stimulation minus baseline) for connections between glutamatergic → glutamatergic **(A)** and glutamatergic → GABAergic **(B)** populations across inter-train intervals (ITIs). The GABAergic → glutamatergic connection was held constant due to limited constraints for reliable parameter estimation in the biophysical model. **(C)** Mean synaptic changes for glutamatergic → glutamatergic connections separated into potentiation and depression components. Two-way ANOVA revealed significant effects of synaptic state (potentiation vs. depression), ITI, and their interaction (p < 0.0001 for all), indicating ITI-dependent modulation of excitatory synaptic plasticity. (D) Mean synaptic changes for glutamatergic-to-GABAe Aergic connections showed similar significant effects (p < 0.0001), with a pronounced bias toward synaptic depression relative to potentiation across ITIs. **(E)** Two-way ANOVA on net synaptic plasticity (potentiation minus depression) revealed a no significant effect connections but significant ITI effect (p<0.0001) and interaction effect (p=0.049). Numerically, there was a dominant negative shift at ITI 20s (eTBS) in both connection types, with a substantially stronger effect in glutamatergic → GABAergic connections compared to glutamatergic → glutamatergic connections, indicating preferential weakening of excitatory drive onto inhibitory neurons. **(F)** Schematic summary illustrating the model-derived mechanism: extended-interval TBS (ITI 20 s) induces modest depression within excitatory recurrent connections while producing robust synaptic depression of glutamatergic inputs onto GABAergic interneurons, resulting in disinhibition of excitatory networks.

To quantify synaptic changes predicted by the model, we analyzed mean weight changes separately for potentiated and depressed synapses. In glutamatergic-to-glutamatergic connections, potentiation and depression differed significantly across ITIs, with a significant interaction between the two (p < 0.0001), indicating that excitatory recurrent plasticity is strongly modulated by stimulation timing (**Fig. 5C**). A similar pattern was observed for glutamatergic-to-GABAergic connections (p < 0.0001), with both synapse types showing a bias toward depression across ITIs, though this effect was considerably more pronounced for glutamatergic-to-GABAergic connections (**Fig. 5D**).

To assess the overall balance of plasticity, we computed net synaptic change (potentiation minus depression) across ITIs. Across conditions, net plasticity was negative for both synapse types, confirming the depression bias observed above. The eTBS with ITI 20s produced the strongest net depression for both connection types, with a substantially larger effect in glutamatergic-to-GABAergic compared to glutamatergic-to-glutamatergic connections (**Fig. 5E**). Together, these modeling results provide a mechanistic framework linking our experimental observations to circuit-level dynamics. It suggests that eTBS modulates excitability both directly (on activated neurons) and indirectly, by depressing excitatory inputs onto GABAergic interneurons, thereby reducing inhibitory tone and enabling sustained glutamatergic activation (**Fig 5F**).

### GABAergic Recruitment Suppresses the Behavioral and Structural Effects of Extended-Interval TBS

The preceding findings showed that eTBS (20s ITI) modulates the excitation / inhibition balance by increasing excitability of glutamatergic neurons while simultaneously weakening glutamatergic inputs onto inhibitory neurons. Moreover, these effects seem to occur via lower recruitment of GABAergic cells during the TBS itself (especially relative to shorter interval TBS paradigms). To test whether these neural effects are relevant to behavior, we first examined both rapid and sustained antidepressant-like outcomes in the CORT model of depression (**Fig 6A)**. We probed effects after a single day of stimulation (rapid antidepressant effects) and after prolonged stimulation (10 sessions), after 21 days of corticosterone exposure. To assess rapid behavioral effects, we performed a single session of stimulation (iTBS-10, eTBS or sham i.e. no stimulation) and compared effects to a group of animals that received saline instead of corticosterone. We first performed an ANOVA across these four groups. and found significant group effect (F (3, 34) = 4.986 p=0.0057). Post-hoc tests were then run to assess group differences. We first confirmed that CORT + sham affected behavior relative to saline (p = 0.011). Next we compared the various CORT groups. Following a single stimulation session, eTBS significantly increased sucrose preference compared to sham-stimulation (p =0.021), indicating a rapid antidepressant-like effect, whereas standard TBS (ITI 10s) did not differ significantly from sham (p =0.64) (**Fig 6B**). After repeated daily stimulation (10 total) we conducted a second behavioral assessment using the forced swim test, a distinct assay chosen to avoid confounds associated with repeated exposure in the sucrose preference test ^21,22^. Our ANOVA analyses indicated significant group effect (F (3, 34) = 11.84, p=<0.0001) and post hoc comparison showed that both iTBS and eTBS groups exhibited significantly reduced immobility times compared to sham (p = 0.0007 and p = 0.0011 respectively; **Fig 6C**), indicating that with repeated stimulation both iTBS and eTBS shows antidepressant-like effects, consistent with data in humans.

**Figure 6.**
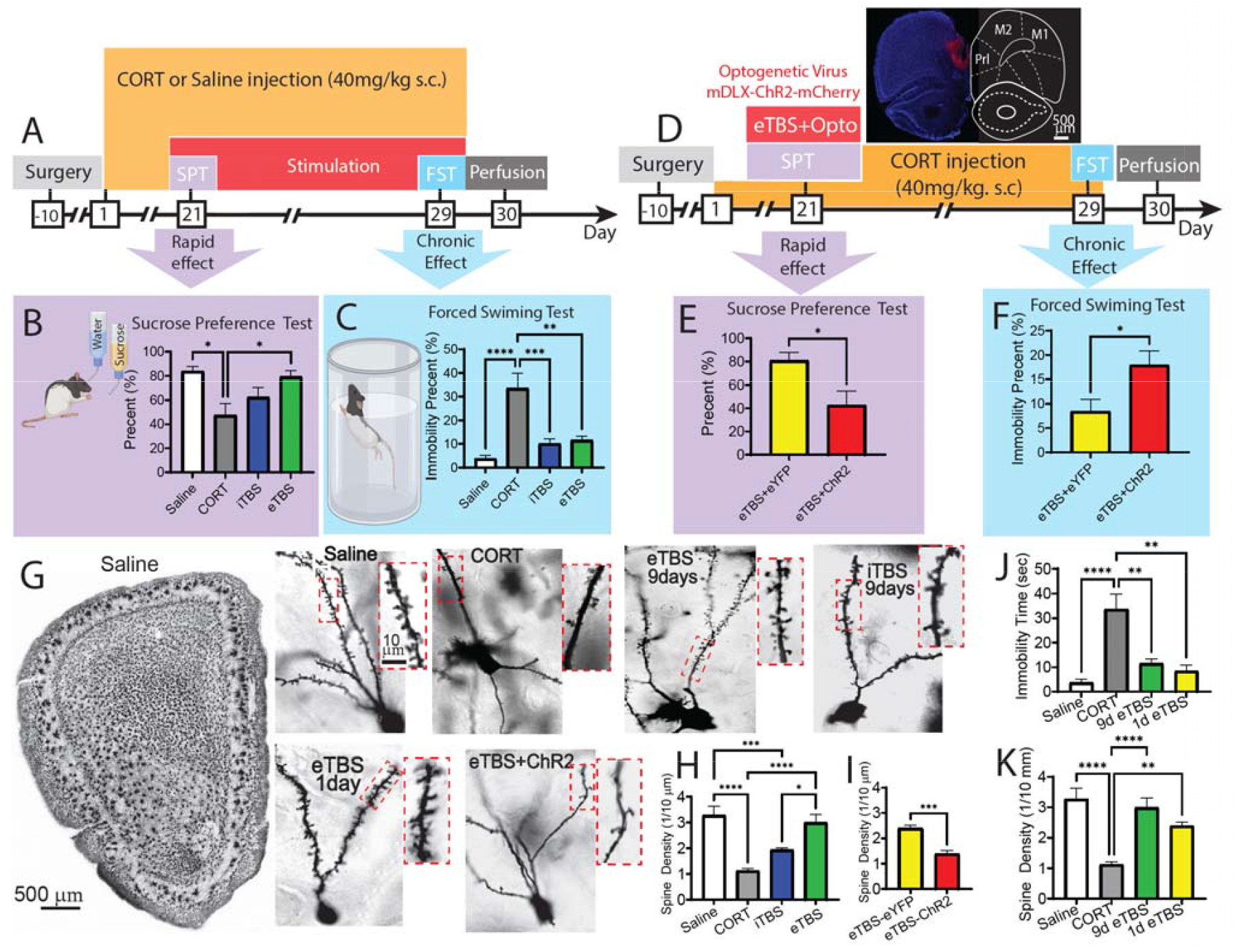
Extended-Interval TBS Induces Antidepressant Effects and Spine Remodeling via Reduced GABAergic Recruitment. **(A)** Experimental timeline. Rats underwent mPFC electrode implantation followed by recovery and daily corticosterone (CORT; 40 mg/kg, s.c.) or saline injections for 3 weeks to induce depression-like behavior. Beginning on day 21, animals received sham, standard TBS (ITI 10 s), or extended-interval TBS (eTBS; ITI 20 s) for 9 consecutive days. **(B)** Sucrose preference test (SPT) following 48-h habituation. CORT-treated animals showed reduced sucrose preference compared to saline controls (p = 0.011), confirming induction of depression-like behavior. A single session of eTBS significantly increased sucrose preference relative to sham (p = 0.021), whereas ITI 10 s stimulation showed no significant effect. **(C)** Forced swim test (FST) following repeated stimulation. Both ITI 10s and ITI 20s protocols significantly reduced immobility time relative to sham (p = 0.0007 and p = 0.0011, respectively), indicating sustained antidepressant-like effects. **(D)** Experimental timeline for optogenetic manipulation. Following viral expression and CORT model induction, animals received a single session of eTBS (ITI 20 s) paired with GABAergic optogenetic stimulation. **(E)** SPT following optogenetic manipulation. Activation of GABAergic neurons during eTBS reduced sucrose preference compared to sham (p = 0.025). **(F)** FST performed 9 days later showed increased immobility time in animals receiving GABAergic optogenetic stimulation during eTBS compared to sham optogenetic controls (p = 0.027), indicating GABAergic recruitment disrupts eTBS anti-depression effect. **(G)** Representative Golgi staining images for dendritic spine analysis. **(H)** Quantification of spine density. CORT significantly reduced spine density (p < 0.0001), whereas eTBS restored spine density relative to CORT-treated animals (p < 0.0001). No significant difference was observed between ITI 10 s stimulation and CORT-treated animals. **(I)** GABAergic controls (p = 0.0003). **(J)** Long lasting anti-depressant comparison between single versus multiple eTBS sessions. Both conditions significantly reduced immobility time relative to CORT-treated animals without stimulation (p = 0.001 for both), with no significant difference between single and repeated sessions, indicating that a single eTBS session produces long-lasting antidepressant-like effects. eTBS-eYFP animals were used as controls for both single-session and optogenetic conditions. **(K)** Spine density comparison between single versus multiple eTBS sessions. Both conditions significantly increased compared to CORT-treated animals without stimulation (p <0.0001 for 9 days eTBS and p=0.0074 for single session eTBS), with no significant difference between single and repeated sessions, indicating that even a single eTBS session produces l restored spine density. eTBS-eYFP animals were used as controls for both single-session and optogenetic conditions. Data were analyzed using one-way ANOVA with Sidak’s post hoc correction and independent t-tests for optogenetic comparisons. Values are mean ± SEM (n = 30 rats, ∼10 per group for behavioral experiments; ∼8 per group for optogenetic and spine density analyses). *p < 0.05, **p < 0.001, ***p < 0.001, ***p < 0.0001.

We performed a follow-up experiment with two goals in mind: 1) do the rapid antidepressant effects observed after a single eTBS last without any repeated stimulation sessions; 2) does increasing GABA activity during eTBS block the behavioral effects. To assess this, we randomized animals into two groups and injected viral constructs encoding either channelrhodopsin (mDLX-ChR2) or a fluorophore control into the mPFC at the site of stimulation. Animals received 21 days of CORT (as above) followed by a single session of eTBS + 20Hz laser stimulation using a custom-designed optrode system described in our previous publication ^23^ (**Fig 6D**). Laser stimulation was delivered starting 200 ms prior to each TBS train and remained on throughout the 2s TBS train but was turned off during the inter-trial period in both groups. Activating the GABAergic neurons blocked the antidepressant effects of eTBS (p = 0.025; **Fig 6E**) and importantly resulted in increased immobility time in the forced swim test measured 8 days later (p = 0.027; **Fig 6F**).

To determine whether these rapid and durable behavioral effects were accompanied by structural plasticity, we performed Golgi staining to assess dendritic spine density across the animals described above (**Fig 6G**). Using a linear mixed-effects model we did not find a significant effect of animal in the model (p = 0.967). Our ANOVA analyses indicated a significant group effect (F(3, 24) = 16.69, p<0.0001) CORT treatment significantly reduced spine density compared with saline controls (p < 0.0001). Prolonged stimulation with eTBS across 10 sessions restored spine density compared with CORT-treated animals (p < 0.0001; **Fig 6H**). In contrast, standard iTBS using a 10 s ITI did not significantly differ from CORT-treated animals, though there was a trend effect (p = 0.15), indicating incomplete structural recovery. Finally, optogenetic activation of GABAergic interneurons during eTBS significantly reduced spine density compared with sham optogenetic controls (p = 0.0003; **Fig 6I**), paralleling the loss of behavioral efficacy and supporting the idea that suppression of excessive inhibitory recruitment is required for eTBS-induced structural restoration.

To better understand the longevity of a single eTBS session, we compared effects on spines/behavior across experiments. Our ANOVA analyses indicated a significant group effect (F (3, 32) = 11.49, p<0.0001) and post hoc comparisons reported that both 1D and 9D eTBS stimulation significantly reduced immobility time measured on day 9 relative to CORT-treated animals without stimulation (p = 0.001 for both), with no significant difference between single and repeated sessions (p = 0.99) or between those conditions and the saline control (p = 0.97 for 1D vs saline, and p = 0.72 for 9D vs saline) (**Fig 6J**). Similarly showed significant group effect (F (3, 24) = 15.10, p<0.0001). Post hoc comparison indicated that both single and repeated eTBS significantly increased spine density compared to CORT-treated animals (p < 0.0001 for 9 days eTBS and p = 0.0074 for single session eTBS), with no significant difference between the two conditions (p = 0.46), or between either condition and saline (p= 0.1 for 1D vs saline and p= 0.96 for 9D vs saline.) (**Fig 6K**). These results indicate that a single session of eTBS is sufficient to produce long-lasting antidepressant-like effects and restore synaptic structure back to nromal. Together, these findings demonstrate that extended-interval TBS induces rapid and durable antidepressant-like effects, accompanies by synaptic plasticity, that depends on reduced GABAergic recruitment.

## Discussion

This study provides evidence that the timing of theta burst stimulation (TBS) trains critically governs excitatory (glutamatergic) and inhibitory (GABAergic) neuron dynamics along with longer-term changes in synaptic plasticity and behavior. Short inter-train intervals (e.g. 4s) induced an immediate surge of calcium activity in both glutamatergic and PV-GABAergic neurons during stimulation, whereas extended interval TBS (20s ITI) emerged as a “sweet spot” that maximized long-term activation and synaptic plasticity of glutamatergic neuron with minimal activation of GABAergic PV cells. Importantly, these circuit-level dynamics translated directly to behavior and synaptic structure in the corticosterone (CORT) model of depression: extended-interval TBS (eTBS) produced rapid and long-lasting antidepressant-like effects, restored dendritic spine density, and, critically, these effects were abolished by optogenetic activation of GABAergic interneurons during stimulation. This causal manipulation demonstrates that reduced inhibitory recruitment is required for both the behavioral and structural efficacy of eTBS. Together, these findings represent the first systematic, single-cell analysis across a range of ITIs in both excitatory and inhibitory cortical neurons, linking stimulation timing to circuit dynamics, behavioral outcomes, and structural plasticity, and revealing how patterned stimulation can shift cortical networks between suppressed and potentiated states.

### Short ITI-TBS Enhances GABAergic / PV interneuron activity limiting longer-term plasticity

Huang et al. (2005) first demonstrated in humans that continuous TBS ( 2s TBS with a 2s ITI, e.g. no pause) suppressed cortical excitability, whereas when TBS with a 10s interval enhanced excitability ^6^. This has since been validated across both human and animal cortex ^6,24–26^. Our results with electrical stimulation in rodents align with these findings and provide novel insight into the mechanisms whereby ITI determines the direction and stability of plasticity following TBS. We first observed that short ITIs strongly increased calcium activity in GABAergic neurons during stimulation, impairing overall synaptic and network plasticity. Prior work (primarily in slice), has shown that excess GABA during TBS stimulation blocks LTP ^27,28^. Pharmacological studies have long shown that blocking GABA receptors permits or amplifies LTP induction ^29,30^. Indeed, it has been shown that if inhibition is triggered during high-frequency stimulation, it can lead to LTD or no potentiation ^17,31^. Thus, our data that short ITI TBS protocols result in preferential activation of GABA inter-neurons provides a clear mechanistic explanation that has been thus far missing for how ITI affects changes in excitability.

Immunohistochemical data showed that fast-spiking PV interneurons appeared to be preferentially driving the increased inhibitory response under short ITI TBS protocols ^18,32^. PV interneurons are well-known for their ability to fire at high frequencies with precise timing and minimal adaptation and target the peri somatic region of pyramidal cells to provide powerful inhibition ^18,27^. Their intrinsic properties (low input resistance, fast membrane kinetics) essentially make them “synchrony detectors” that preferentially respond to the kind of tightly clustered input that short-interval TBS provides ^33^. Once recruited, PV neurons can rapidly release GABA and strongly curtail pyramidal cell spiking. Prior literature supports this view; PV interneurons have been implicated in controlling the gain and timing of network excitation, effectively creating a “ceiling” on pyramidal cell activity during intense stimulation ^18,27^. PV neurons have been implicated in the response to rTMS in a wide range of animal models, with evidence for local and network wide effects ^34,35 27^.

### Is there an optimal ITI for plasticity induction with TBS?

Prior work in humans examining the physiological effects of iTBS (10s ITI), has shown highly variable effects. While there is a clear “mean” effect on increasing excitability ^4,6,10^, there is a high degree of variability in these effects ^36–39^. This variability has been posited to limit the clinical efficacy of TBS paradigms. Our data indicates that the standard 10s ITI is not optimized for inducing cortical plasticity, providing a mechanistic basis for the modest and variable physiological effects commonly observed in human and clinical studies. The U-shaped relationship we observed implies that two competing processes are at play. At very short ITIs, inhibitory processes likely accumulate and restrict plasticity. However, at extremely long ITIs (60s), each burst train may act in isolation, failing to build up the postsynaptic calcium or kinase signaling cascades needed for sustained LTP ^4,40,41^. Differential analyses reveal a clear “sweet spot” at an ITI of 20 s, where excitatory neurons reach maximal enhancement while inhibitory neurons exhibit maximal suppression (Fig 6D). The sustained glutamatergic activation observed at this interval likely reflects an optimal excitation-inhibition balance, in which repeated depolarizing inputs are not prematurely curtailed by inhibitory activity, thereby enabling robust synaptic strengthening. By comparison, at ITI-10s (the FDA-cleared standard), concurrent activation of excitatory and inhibitory populations appears to counteract one another, effectively dampening net plasticity and contributing to the variability reported in human studies. Under these conditions, individual differences in baseline excitation-inhibition balance are likely to exert a dominant influence on stimulation outcomes. Consistent with this interpretation, baseline cortical excitability has been identified as a key predictor of response to iTBS (ITI 10s) in both healthy individuals and patients with depression.^42–44^.

### eTBS Effects on Cortical Disinhibition and Synaptic Plasticity

Intermittent theta-burst stimulation has been shown to trigger lasting decreases in GABA release / interneuron firing, mediated by mechanisms such as GABA-B auto receptor activation or long-term depression at interneuron synapses ^17,27,30^. We observed, particularly for extended TBS paradigms, that it induces a lasting reduction in their post-stimulation activity, an effect not readily driven by an increase in calcium levels. Our RNN model, trained directly on empirical data, revealed that the network level changes observed with ITI 20s are mediated by a selective depression of glutamatergic-to-GABAergic synapses that outpaces depression of glutamatergic-to-glutamatergic connections - an effect that is most pronounced at ITI 20s and shifts the overall excitation-inhibition balance toward long-term disinhibition, a weakening of inhibitory tone that would favor increased network excitability ^45,46^.

Importantly, our finding that a single session of eTBS is sufficient to produce both long-lasting antidepressant-like effects and sustained increases in dendritic spine density provides key insight into the temporal dynamics of TBS-induced plasticity. These results suggest that extended-interval stimulation engages rapid mechanisms capable of triggering enduring circuit reorganization, rather than requiring cumulative effects across repeated sessions. One plausible interpretation is that the reduced recruitment of GABAergic interneurons during eTBS lowers the threshold for activity-dependent synaptic strengthening, enabling a single stimulation bout to initiate cascades associated with long-term potentiation and structural remodeling ^47–50^. This interpretation is consistent with extensive evidence that disinhibition facilitates synaptic plasticity by permitting sufficient postsynaptic depolarization and calcium influx to engage downstream signaling pathways ^29,51^. In contrast, excessive inhibitory recruitment, as observed at shorter ITIs or during optogenetic activation of GABAergic neurons, may prevent these processes by constraining excitatory activity and limiting structural plasticity. The persistence of both behavioral and spine changes following a single eTBS session further suggests that extended ITIs engage mechanisms resembling metaplasticity or consolidation processes that stabilize newly formed synaptic states^52,53^. Together, these findings highlight that the temporal structure of stimulation not only determines the immediate excitatory-inhibitory balance but also governs the induction and stabilization of long-term structural and behavioral plasticity.

## Conclusion

Our results collectively indicate that the stability of neuronal responses following extended ITIs arises from a coordinated rebalancing of glutamatergic and GABAergic activity. Short ITIs, despite eliciting strong immediate responses, disrupt long-term network stability, likely due to excessive inhibitory recruitment and impaired excitation-inhibition balance. In contrast, moderate ITIs, particularly around 20 seconds, promote an optimal interaction between excitatory and inhibitory populations, enabling sustained post-stimulation excitability. Critically, this circuit-level mechanism translates to behavior and synaptic structure: extended-interval TBS produces rapid and durable antidepressant-like effects in the CORT model, restores dendritic spine density, and these benefits are abolished when GABAergic interneurons are artificially recruited during stimulation, demonstrating a causal role for disinhibition. These findings establish temporal spacing as a key determinant of enduring cortical plasticity and stability. More broadly, they provide a mechanistic framework for reducing variability in current iTBS protocols and suggest that extending the inter-train interval may enhance the efficacy and reliability of neuromodulation therapies by maximizing sustained excitatory responses while limiting inhibitory overactivation.

## Supporting information

Figures an Tables for further stats

## Resource Availability

Further information and requests for resources should be directed to and will be fulfilled by the Lead Contact, Dhakshin Ramanathan and Morteza Salimi

## Materials and Methods

### Subjects

A total of 90 female Long Evans rats were used across all experiments. Rats were obtained from Charles River Laboratories and arrived at approximately 150 g and one month of age. Animals were housed in pairs in standard plastic cages during acclimation and were allowed to acclimate for two weeks before surgery. Following surgery, rats were individually housed to facilitate recovery. Animals were maintained on a 12 h light/dark cycle, with lights on at 6:00 a.m., and all experimental procedures were performed during the light phase. Food and water were available ad libitum. At the time of recording or behavioral testing, rats weighed approximately 300–500 g and were 3–5 months old.

For the initial in vivo calcium imaging experiments, 10 rats were used, including 5 rats for glutamatergic imaging and 5 rats for GABAergic imaging. An additional 8 rats were excluded from imaging analyses because of electrolens misplacement. Histological analyses were performed in 20 rats (5 animals per group), with 3 additional rats excluded because of staining failure. Behavioral experiments were conducted in 38 rats, and optogenetic experiments were conducted in 14 rats. Remaining animals were used for validation, surgical optimization, and replacement of excluded animals across experiments.

### Electrolens Fabrication

To deliver theta burst stimulation (TBS), electrodes were fabricated by attaching four 50 μm tungsten wires (California Fine Wire) with epoxy alongside an integrated ProView lens. Wires were spaced 100 μm apart beneath the lens edge. Optimal electrodes for effective calcium response were selected. A 75 μm stainless-steel wire (A-M Systems, Sequim, WA, USA) served as the ground, soldered to a stainless steel bone screw (Fine Science Tools).

### Surgical Procedures

Stereotaxic surgery was performed under sterile conditions. Rats were anesthetized initially with 5% isoflurane and maintained at 1.9–2.5% throughout surgery, with body temperature regulated via a heating mat (VWR, Radnor, PA, USA). Atropine (0.05 mg/kg) reduced respiratory secretions; Dexamethasone (0.5 mg/kg) minimized inflammation; saline (1 mL, 0.9%) provided hydration; and Lidocaine (0.2 cc max) offered local analgesia. After incision and skull exposure, anchor and ground screws were inserted. A hole (1.2 mm diameter) was drilled at coordinates targeting the medial prefrontal cortex (mPFC; AP=+4.2 mm, ML=0.7 mm). Viral vectors (targeting either glutamatergic [CamKII-driven GCaMP] or GABAergic [mDlx-driven GCaMP] neurons) were infused (100 nL/min) at DV=2.8 mm and allowed to diffuse for 10 min post-infusion. An integrated ProView lens (1 mm diameter, 9 mm length) was implanted at DV=2.4 mm, sealed with Kwik-Sil (WPI), and secured using dental cement. Electrodes were connected to a TDT ZCA-EIB32 ZIF-Clip® Headstage-to-Electrode Interface Board. Post-surgical analgesics were administered as required. Procedures adhered strictly to NIH guidelines and were approved by the San Diego VA Medical Center IACUC (Protocol Number A12-021).

### Data Processing and Analysis

In vivo imaging captured cellular fluorescence during and for 40 minutes post-TBS. Data were motion corrected, segmented, and ΔF/F traces extracted using CNMF-E. Neurons were classified as excited, inhibited, or unchanged based on z-scored fluorescence during stimulation and post-stimulation epochs. Data processing and analysis were conducted using MATLAB v2023a. Cells were classified based on their normalized activity throughout the entire recording session. Normalization was performed using Z-score transformation, with the 5-minute period preceding the onset of TBS serving as the baseline for each cell. Cells were classified as ‘excited’, ‘inhibited’, or ‘non-responsive’ based on their activity during the TBS period compared to baseline activity. To reduce noise and ensure cleaner comparisons, average activity within 4-second time bins was used, resulting in 15 samples per minute of recording. Classification was carried out using the Wilcoxon rank-sum test in MATLAB, with a significance threshold of p < 0.01.

Additionally, for analyses involving neuronal response stability (Figure 4), cells were further sorted based on their activity across the entire 40-minute post-stimulation period relative to baseline, allowing for evaluation of the consistency of neuronal identity over time.

### Theta Burst Stimulation (TBS) Protocol

TBS was administered via a Tucker-Davis Technologies IZ2H stimulator and passive ZIF-Clip® headstages. Electrical stimulation was comprised of 20 trains of 10 bursts, with each burst containing 3 biphasic, cathode-leading pulses (100 µA, 400 µs pulse width). Inter-train intervals (ITIs) ranged from 4 to 60 seconds, defined from the start of consecutive trains. Bursts and pulses were administered at 5 Hz and 50 Hz, respectively. The stimulation was facilitated via tungsten electrodes that were secured to the imaging lens.

### Sham Experimental Condition

For the calcium imaging experiments, animals previously used in stimulation sessions were reconnected to the imaging and stimulation apparatus under identical conditions in the separate session, but no current was delivered. Data from these sessions were processed using the same analytical pipeline as the stimulation experiments to ensure methodological consistency. In the histological experiments, sham animals underwent identical surgical implantation of electrodes and were connected to the stimulation device without receiving any stimulation, thereby controlling for procedural and handling variables.

### Histological Assessment

#### Tissue Preparation

At the conclusion of calcium imaging experiments, rats were deeply anesthetized and transcardially perfused with 0.9% saline followed by 4% paraformaldehyde (PFA) in phosphate-buffered saline (PBS). Extracted brains were post-fixed overnight at 4°C in 4% PFA, followed by cryoprotection in 30% sucrose solution in PBS until fully submerged. Coronal brain sections (40 µm thickness) were prepared using a cryostat (Leica CM1950, Leica Biosystems) and collected onto gelatin-coated slides. Sections were air-dried, rehydrated in PBS, and stored at 4°C until processing.

#### Verification of Viral Expression in Calcium Imaging Rats

To confirm successful viral delivery and expression in rats utilized for calcium imaging, brain sections were mounted onto slides, counterstained with DAPI (4′,6-diamidino-2-phenylindole), and coverslipped with antifade mounting medium (Fluoromount-G). Images were captured using confocal microscopy (Zeiss LSM 880, Zeiss Microscopy) and matched to the Paxinos and Watson rat brain atlas to verify accurate targeting of the medial prefrontal cortex.

#### Immunohistochemical Assessment of c-Fos and GAD-65

A separate cohort of rats (n = 4 per group) underwent TBS at ITIs of 4, 10, or 20 seconds or served as sham (electrode implantation and connection without stimulation), and were perfused 60 minutes post-stimulation. Sections underwent either double or triple immunofluorescence staining.

#### Double Staining (c-Fos and GAD-65)

One section per animal was incubated overnight at 4°C with primary antibodies purchased from Abcam:

- c-Fos: Anti-c-Fos monoclonal antibody [2H2], host: Mouse (Abcam, cat# ab208942, 1:1000 dilution)
- GAD-65: Anti-GAD-65 monoclonal antibody [EPR22952-70], host: Rabbit (Abcam, cat# EPR22952-70, 1:300 dilution)

Sections were subsequently incubated for 2 hours at room temperature with secondary antibodies from Thermo Fisher (1:500 dilution):

- Donkey anti-Mouse IgG (H+L), Alexa Fluor™ Plus 488 (Thermo Fisher, cat# A32766, green)
- Donkey anti-Rabbit IgG (H+L), Alexa Fluor™ 568 (Thermo Fisher, cat# A10042, red)

Quantification of immunoreactivity intensity for double staining was conducted using FIJI (ImageJ). Regions of interest (ROIs; 500 µm spanning around the stimulation electrode) were analyzed and data averaged per animal for statistical comparisons.

#### Triple Staining (PV, SST, and GAD-65)

Another section per animal was incubated overnight at 4°C with primary antibodies purchased from Abcam:

- Parvalbumin (PV): Polyclonal antibody, host: Chicken (Abcam, cat# PA5-143579, 1:300 dilution)
- Somatostatin (SST): Monoclonal antibody [7G5], host: Mouse (Abcam, cat# MA5-17182, 1:300 dilution)
- GAD-65: Anti-GAD-65 monoclonal antibody [EPR22952-70], host: Rabbit (Abcam, cat# EPR22952-70, 1:300 dilution)

Sections were subsequently incubated for 2 hours at room temperature with secondary antibodies from Thermo Fisher (1:500 dilution):

- Donkey anti-Chicken IgY (H+L), Alexa Fluor™ 647 (Thermo Fisher, cat# A78952, far red, PV)
- Donkey anti-Mouse IgG (H+L), Alexa Fluor™ Plus 488 (Thermo Fisher, cat# A32766, green, SST)
- Donkey anti-Rabbit IgG (H+L), Alexa Fluor™ 568 (Thermo Fisher, cat# A10042, red, GAD-65)

Confocal microscopy (Zeiss LSM 880) was used for high-resolution image acquisition. For quantitative analyses, Imaris software (Oxford Instruments) was employed to perform three-dimensional cellular reconstruction, quantify immunoreactivity intensity within individual cells, and assess colocalization of GAD-65 with PV and SST interneurons using Mander’s coefficients only in ROI-1 corresponding 500 µm spanning around the stimulation electrode.

#### Golgi staining and spine density analysis

To assess structural plasticity, dendritic spine density was quantified using the FD Rapid GolgiStain™ Kit (FD NeuroTechnologies) according to the manufacturer’s instructions. Following staining, coronal brain sections containing the medial prefrontal cortex (mPFC) were imaged using a Keyence fluorescent microscope equipped with a 100× objective lens. Spine density analyses were performed on clearly isolated dendritic segments from stained pyramidal neurons within the prelimbic mPFC. Z-stack images were imported into Imaris software (Oxford Instruments), and dendritic reconstructions were generated using the Filament Tracer module. Dendritic shafts were semi-automatically traced, and spines were detected and manually verified to ensure accurate segmentation. Spine density was calculated as the number of spines normalized to dendritic length (spines/µm) and averaged across segments for each animal prior to statistical analysis.

### Chronic corticosterone model and behavioral testing

#### Corticosterone treatment

To induce depression-like behavior, rats were treated with chronic corticosterone (CORT) injections using a widely adopted high-dose regimen. CORT (40 mg/kg, s.c.; Sigma) was dissolved in a vehicle containing Tween-20, dimethyl sulfoxide (DMSO), and sterile 0.9% saline (adapted from prior CORT vehicle formulations). Injections were administered once daily for three consecutive weeks, resulting in approximately four weeks of continuous CORT exposure when including the subsequent stimulation and behavioral testing period. After CORT treatment, animals were assigned to sham (n = 11), standard TBS (ITI 10s; n = 9), or extended-interval TBS (eTBS; ITI 20s; n = 10) conditions for behavioral experiments.

#### Sucrose Preference Test (SPT) acute antidepressant-like effects

To assess rapid changes in anhedonia-like behavior, we used the sucrose preference test after the first stimulation session. The SPT was performed in CORT-treated rats from the sham, ITI 10 s, and ITI 20 s groups. Forty-eight hours before the main test day, animals underwent a 3h habituation period with access to two identical drinking bottles, one containing water and one containing a 1% sucrose solution. On the test day, rats again received access to two counterbalanced bottles (water vs. sucrose), and fluid intake from each bottle was measured. Sucrose preference was calculated as:

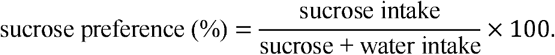

Due to the known sensitivity of SPT outcomes to model duration, deprivation, and test repetition, the SPT was used only to assess the acute behavioral response to a single stimulation session, not for longitudinal tracking.

#### Forced Swim Test (FST) for chronic antidepressant-like effects

Sustained antidepressant-like effects were evaluated using the forced swim test following nine consecutive days of daily TBS or sham stimulation. Rats were individually placed in a transparent cylinder (approx. 50 cm height, 20-22 cm diameter) filled with water (25 ± 1 °C, depth ∼30–35 cm) for a 6 min test session. Behavior was recorded with a top-or side-mounted camera.

Videos were analyzed offline using DeepLabCut for pose estimation of body coordinates, and immobility was quantified with a custom MATLAB script. Immobility was defined as periods in which the animal displayed only minimal movements necessary to keep the head above water, with no active swimming or climbing, in line with standard FST definitions. The MATLAB algorithm used frame-by-frame body velocity and posture to classify behavior, and immobility bouts shorter than ∼1 s were excluded, consistent with threshold-based automated FST scoring approaches. Total immobility time during the 6 min session was used as the primary readout of depression-like behavior.

### Statistical Analysis

Statistical analyses were conducted using IBM SPSS Statistics, Version 31, and GraphPad Prism, Version 10.6, was used for data visualization. Adobe Illustrator 2025 was used for final figure assembly. For single-cell calcium analyses, z-scored calcium activity was used as the dependent variable. To account for potential animal-level variability, we first applied linear mixed-effects models with animal identity included as a random effect, while stimulation condition, inter-train interval (ITI), and the stimulation condition × ITI interaction were included as fixed effects. When animal-level random effects were not significant, follow-up ANOVA models were used to evaluate stimulation-dependent differences across ITIs, with Sidak-corrected within-ITI comparisons used to compare stimulation with ITI-matched sham conditions. For post-stimulation analyses, activity was analyzed across eight discrete 5-min bins spanning the 40-min post-TBS period. For categorical cell-state analyses, cells were classified as activated, inactivated, or neutral based on whether calcium activity significantly increased, decreased, or did not significantly differ from baseline. Cell category data were transformed into binary variables, such as activated versus not activated or inactivated versus not inactivated, to compare proportions between stimulation and sham conditions using Chi-square tests. For transition analyses, activated-during cells were further classified as stable activated if they remained activated post-stimulation, or unstable activated if they transitioned to either neutral or inactivated states after stimulation. Chi-square tests were used to compare the proportion of stable versus unstable activated cells between stimulation and sham conditions within each ITI. All statistical tests were two-tailed, significance was set at p < 0.05, and data are presented as mean ± SEM.

## Acknowledgments

This material is the result of work supported with resources and the use of facilities at the VA San Diego Medical Center and the Center of Excellence for Stress and Mental Health. The contents of this manuscript do not represent the views of the U.S. Department of Veteran Affairs or the United States Government. This study was supported by the following grants. Dhakshin Ramanathan, Burroughs Wellcome Fund (https://dx.doi.org/10.13039/100000861), Award ID: 1015644. Dhakshin Ramanathan, National Institute of Mental Health (https://dx.doi.org/10.13039/100000025), Award ID: R01MH123650. Jyoti Mishra, Hope for Depression Research Foundation. Grant ID: 30063773. Maxim Bazhenov: National Institute of Health, Award ID: 1R01MH125557, 1RF1NS132913, 1R01AG099626 and National Science Foundation, Award ID: BCS-2323241. Dhakshin Ramanathan, UCSD Shiley-Marcos Alzheimer’s Disease Research Center Grant ID:P30 AG062429.We also acknowledge the UCSD School of Medicine Microscopy Core Grant P30 NS047101.The authors and do not necessarily reflect the position or policy of the VA, NIH and other grants. The funders had no role in the design of the study, data analysis, or decision to publish. The authors have no relevant biomedical conflicts of interest to disclose.

## Author contributions

Conceptualization: MS, DR. Technical development: MS, AL, MN. Setting up the stimulation and synchronization: Jonathan M. Investigation: MS. and AL. Analysis: MS and MN, ShJ, RG, MB, JR. Resources and funding acquisition: DR, Jyoti M. Writing original draft: MS, DR, MN SJ, ShJ, and MK; writing reviewing & editing, MS, DR, MN, JR, MB and Jyoti M.

## Declaration of Interest

The author declares no conflicts of interest related to this study. The views expressed in this article are those of the authors and do not necessarily reflect the position or policy of the VA or the NIH. The funders had no role in the design of the study, data analysis, or decision to publish. The authors have no relevant biomedical conflicts of interest to disclose.

## Notes

### Competing Interest Statement

The authors have declared no competing interest.

### Summary of Updates

This revised version includes substantial new analyses and experimental data. We added computational modeling to estimate stimulation-induced changes in effective synaptic connectivity and to support a disinhibitory mechanism underlying extended-interval TBS. We also added new behavioral experiments showing rapid and durable antidepressant-like effects, including optogenetic manipulation demonstrating that GABAergic recruitment suppresses these effects. In addition, we included new histological and spine-density analyses linking extended-interval TBS to reduced inhibitory recruitment and structural plasticity. Figures, supplementary figures, methods, and the discussion were updated accordingly.

