## Supplementary material for "Extended-Interval Theta Burst Stimulation Enhances Glutamatergic Plasticity and Antidepressant Effects via Reduced GABAergic Recruitment": Figures an Tables for further stats

**Materials and Methods**

**Virus Injection and Miniscope Implantation**

To label excitatory or inhibitory neurons, rats were injected with AAV-Syn-GCaMP for glutamatergic neuron imaging or AAV-Dlx-GCaMP for GABAergic imaging. A GRIN lens was implanted above the medial prefrontal cortex for calcium imaging using a Inscopix miniaturized one-photon microscope.

**Theta Burst Stimulation Protocols**

TBS consisted of 20 trains of 2-second bursts (3 pulses at 50Hz repeated at 5Hz) delivered via implanted electrodes targeting the mPFC. Inter-train intervals (ITI) were varied systematically across five conditions (4s, 10s, 20s, 40s, 60s). Sham animals were connected to the stimulator but received no stimulation.

**Calcium Imaging and Data Processing**

In vivo imaging captured cellular fluorescence during and for 40 minutes post-TBS. Data were motion corrected, segmented, and ΔF/F traces extracted using CNMF-E. Neurons were classified as excited, inhibited, or unchanged based on z-scored fluorescence during stimulation and post-stimulation epochs.

**Histology**
Brains were perfused and sectioned for immunohistochemistry. Antibodies against GAD65, PV, and SST were used to assess GABAergic marker expression in the mPFC.

**Statistical Analysis**

Statistical analyses included Chi-square tests, two-way ANOVAs, and McNemar’s tests, with Sidak corrections for multiple comparisons. Data was analyzed using MATLAB and GraphPad Prism.

**Subjects**

The study utilized five female Long Evans rats, obtained from Charles River Laboratories. Upon arrival, rats (~150 g, approximately one-month-old) were housed in pairs within standard plastic cages (Allentown, NJ, USA) and acclimated for two weeks prior to surgery. Post-surgery, animals were individually housed to facilitate recovery. Rats were maintained under a 12-hour light/dark cycle (lights on at 6 a.m.), with experimental procedures conducted during the light phase. Food and water were available ad libitum. At the onset of recording experiments, rats weighed between 300–600 grams and were 3–5 months old. 26 rats included ( 5 per glutamatergic and GABAergic calcium imaging group, 16 rats for histology). 11 rats were excluded for electrolens misplacements (8 rats), and staining failure (3 rats).

To evaluate the relationship between neuronal activity during the TBS and post-TBS periods, a linear regression model was implemented in MATLAB, with activity during TBS as the independent variable (X) and post-TBS activity as the dependent variable (Y). The results included regression coefficients, 95% confidence intervals, R² values (goodness of fit), and Pearson's correlation coefficients, providing insights into the stability and consistency of neuronal responses across different stimulation intervals.

$$\text{sucrose preference (\%)}=\frac{\text{sucrose intake}}{\text{sucrose + water intake}}\times100.$$

Due to the known sensitivity of SPT outcomes to model duration, deprivation, and test repetition, the SPT was used only to assess the acute behavioral response to a single stimulation session, not for longitudinal tracking.

**Supplementary Figures and tables**

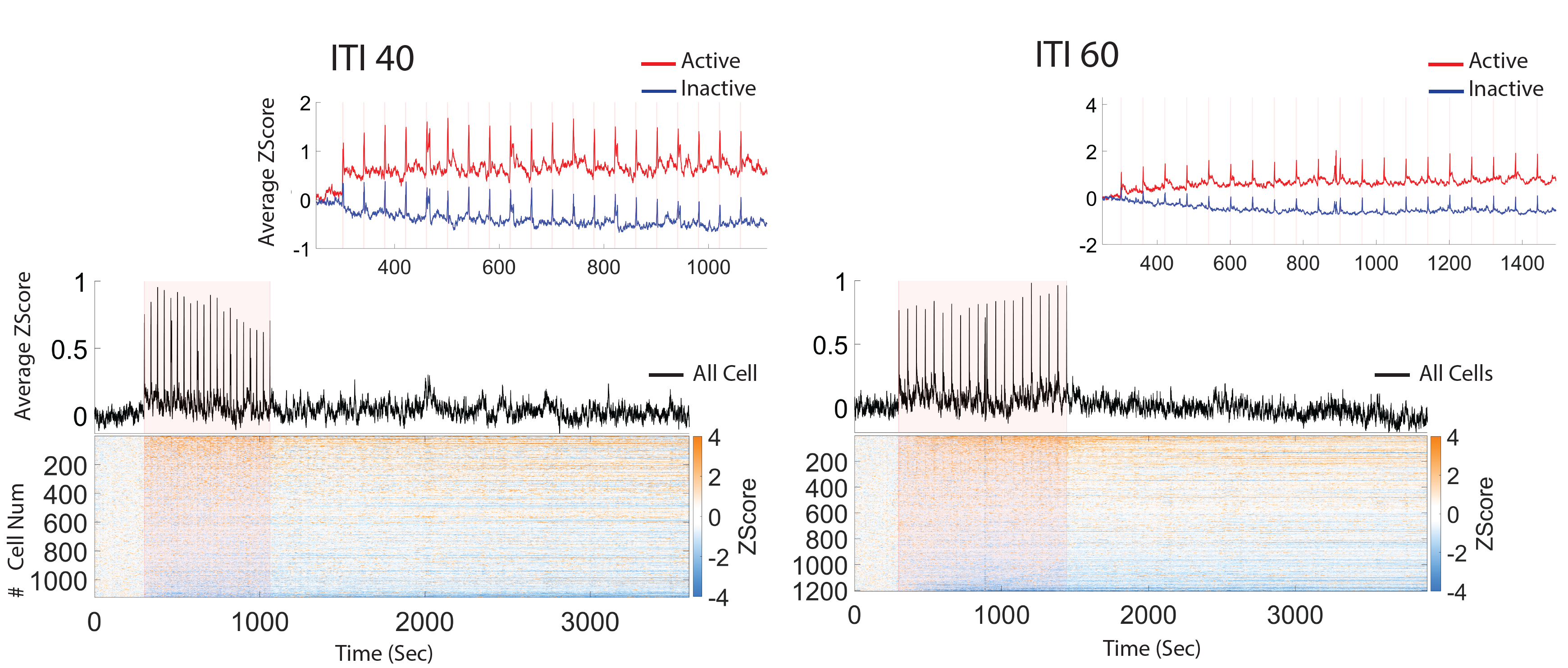
**Figure S1.** Z-scored calcium activity across the full recording session among all recorded cells, ranked from most active (red) to most inactive (blue).

**Table S1. Comparison of Z-scored calcium activity during stimulation across inter-train intervals (ITIs).** Statistical analyses correspond to Figures 1F and 2B. For each ITI (4, 10, 20, 40, and 60 s), Z-scored calcium activity during stimulation was compared between sham and stimulation conditions using two-way ANOVA. Analyses were performed separately for all recorded glutamatergic neurons, activated neurons, and inactivated neurons. Reported values indicate the main effect of stimulation condition (sham vs. stimulation) within each ITI. Activated and inactivated neurons were classified based on significant deviations from baseline calcium activity during stimulation (p < 0.01).

|  | Cell Category in response to stimulation | | | |
| --- | --- | --- | --- | --- |
| ITI (Sec) |  | All | Activated | Inactivated |
|  | 4 | F=570.7  p<0.0001 | F=634  p<0.0001 | F=382.4  p<0.0001 |
|  | 10 | F=70.22  p<0.0001 | F=99.17  p<0.0001 | F=135.8  p<0.0001 |
|  | 20 | F=16.97  p<0.0001 | F=21.39  p<0.0001 | F=31.9  p<0.0001 |
|  | 40 | F=5.23  p=0.022 | F=7.34  p=0.007 | F=0.03  p=0.85 |
|  | 60 | F=2.7  p=0.1 | F=4.81  p=0.028 | F=8.92  p=0.003 |

**Table S2. Comparison of Z-scored calcium activity post-stimulation in glutamatergic neurons across inter-train intervals (ITIs).** Statistical analyses correspond to Figures 1G and 2D. Post-stimulation calcium activity was quantified across eight consecutive 5-min bins spanning 40 min following TBS. For each ITI (4, 10, 20, 40, and 60 s), Z-scored calcium activity was compared between sham and stimulation conditions using two-way ANOVA. Analyses were performed separately for all recorded glutamatergic neurons, activated neurons, and inactivated neurons. Reported statistics reflect the main effect of stimulation condition (sham vs. stimulation) within each ITI. Activated and inactivated neurons were classified based on significant deviations from baseline calcium activity during stimulation (p < 0.01).

|  | Cell Category in response to stimulation | | | |
| --- | --- | --- | --- | --- |
| ITI (Sec) |  | All | Active | Inactive |
|  | 4 | F=35.22  p<0.0001 | F=82.42  p<0.0001 | F=0.4  p=0.5 |
|  | 10 | F=7.16  p=0.007 | F=6.27  p=0.012 | F=0.15  p=0.693 |
|  | 20 | F=17.64  p<0.0001 | F=16.64  p<0.0001 | F=7.64  p=0.006 |
|  | 40 | F=8.35  p=0.004 | F=4.72  p=0.03 | F=4.71  p=0.03 |
|  | 60 | F=0.031  p=0.86 | F=14.21  p<0.0001 | F=11.62  p<0.0001 |

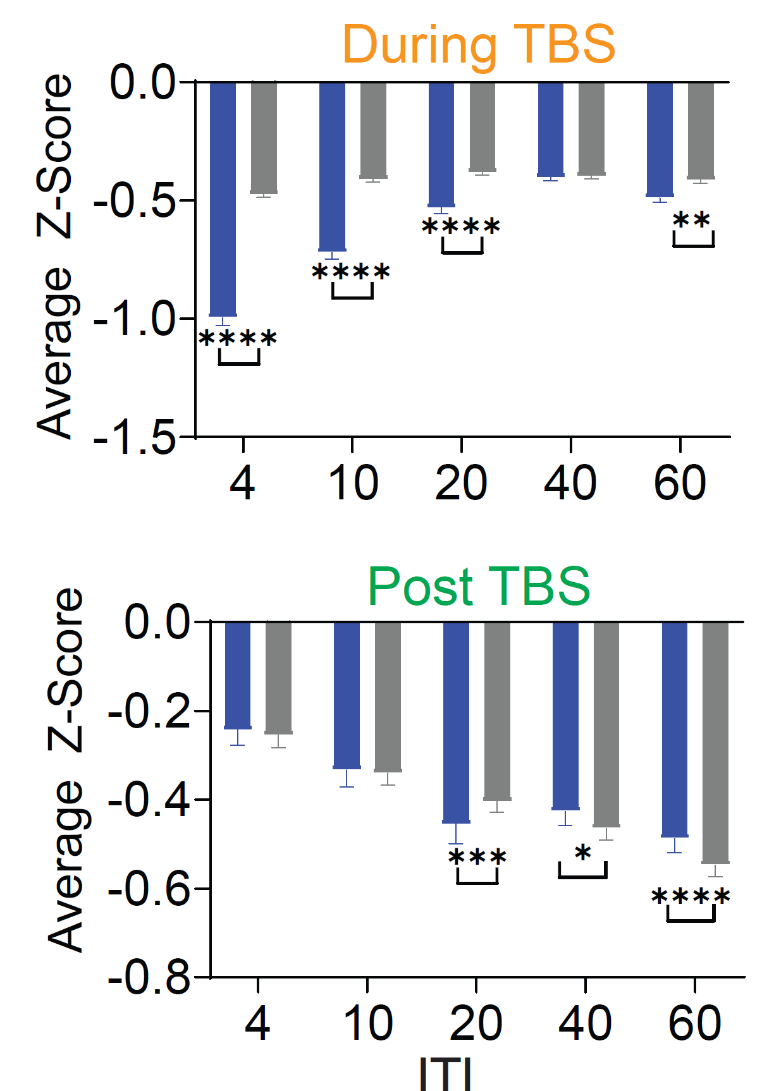

**Figure S2 . Inactivated glutamatergic neurons.** Top panels show calcium activity during TBS, whereas bottom panels show post-stimulation activity. Average calcium activity of inactivated cells was compared between sham and stimulation conditions during and after TBS. During stimulation, inactivated cells exhibited significantly lower calcium activity than sham at ITIs 4, 10, 20, and 60 s, with the strongest suppression observed at shorter ITIs and progressively weaker effects at longer intervals. No significant difference was observed at ITI 40 s. Post-TBS, inactivated cells showed significantly lower activity than sham at ITIs 20, and higher than sham at 40, and 60 s, whereas no significant differences were detected at ITIs 4 and 10 s, indicating that extended ITIs promoted sustained suppression of inactivated neuronal populations after stimulation. Data were analyzed using two-way ANOVA followed by Sidak’s correction for multiple comparisons. Gray indicating sham and blue indicating stimulation condition. Values are presented as mean ± SEM

**Figure S3 . Comparison of Z-scored calcium activity post stimulation.** Post-TBS Z-scored activity 8 discrete of 5min for 40 min post time bins for all, active and inactive cells population. Each point indicates average of Z-score within 5 min time bin. shown mean ±SEM in corresponding fig 2 D. Z-scored calcium activity was calculated by normalizing activity across the session to this baseline.

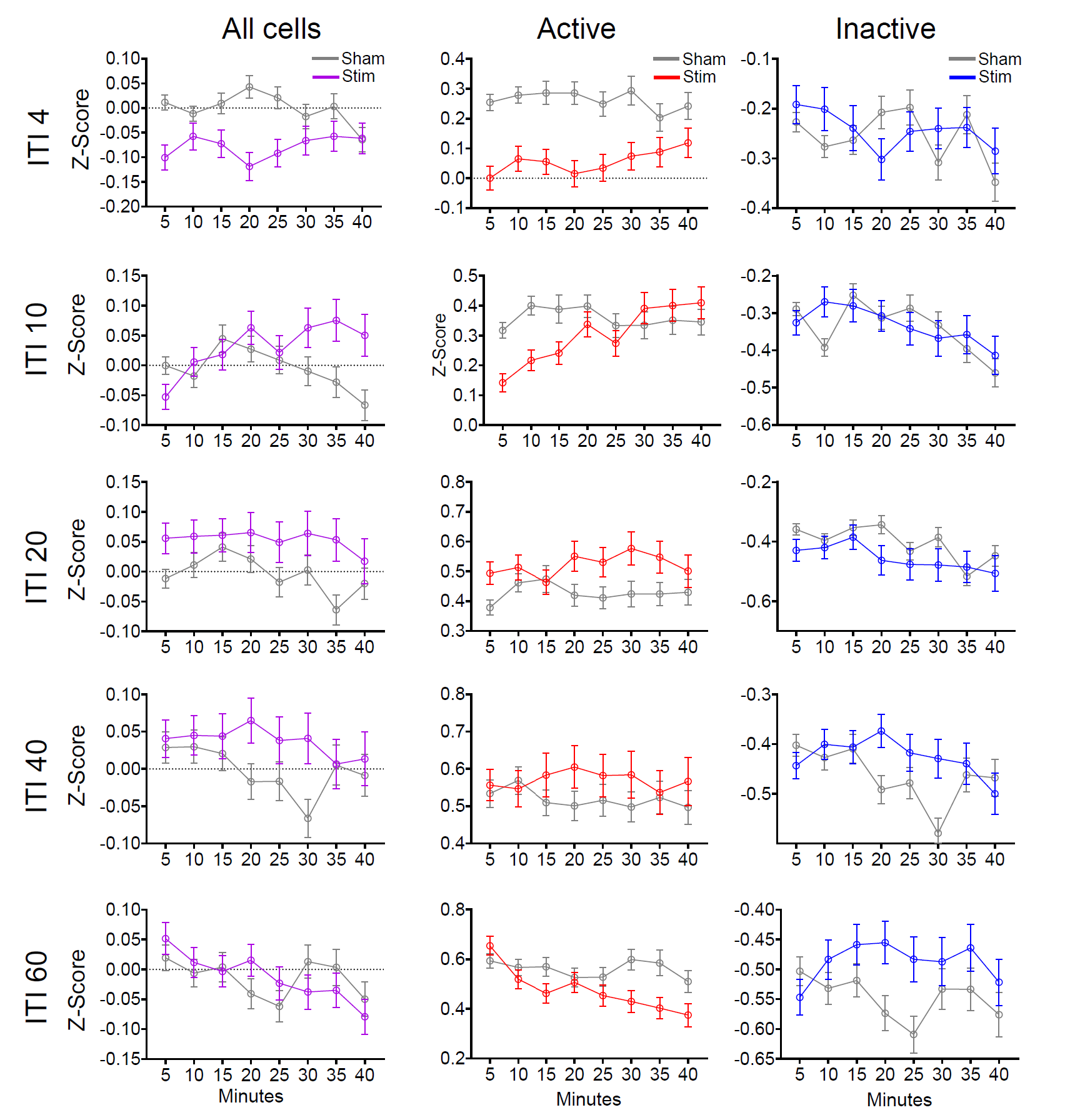

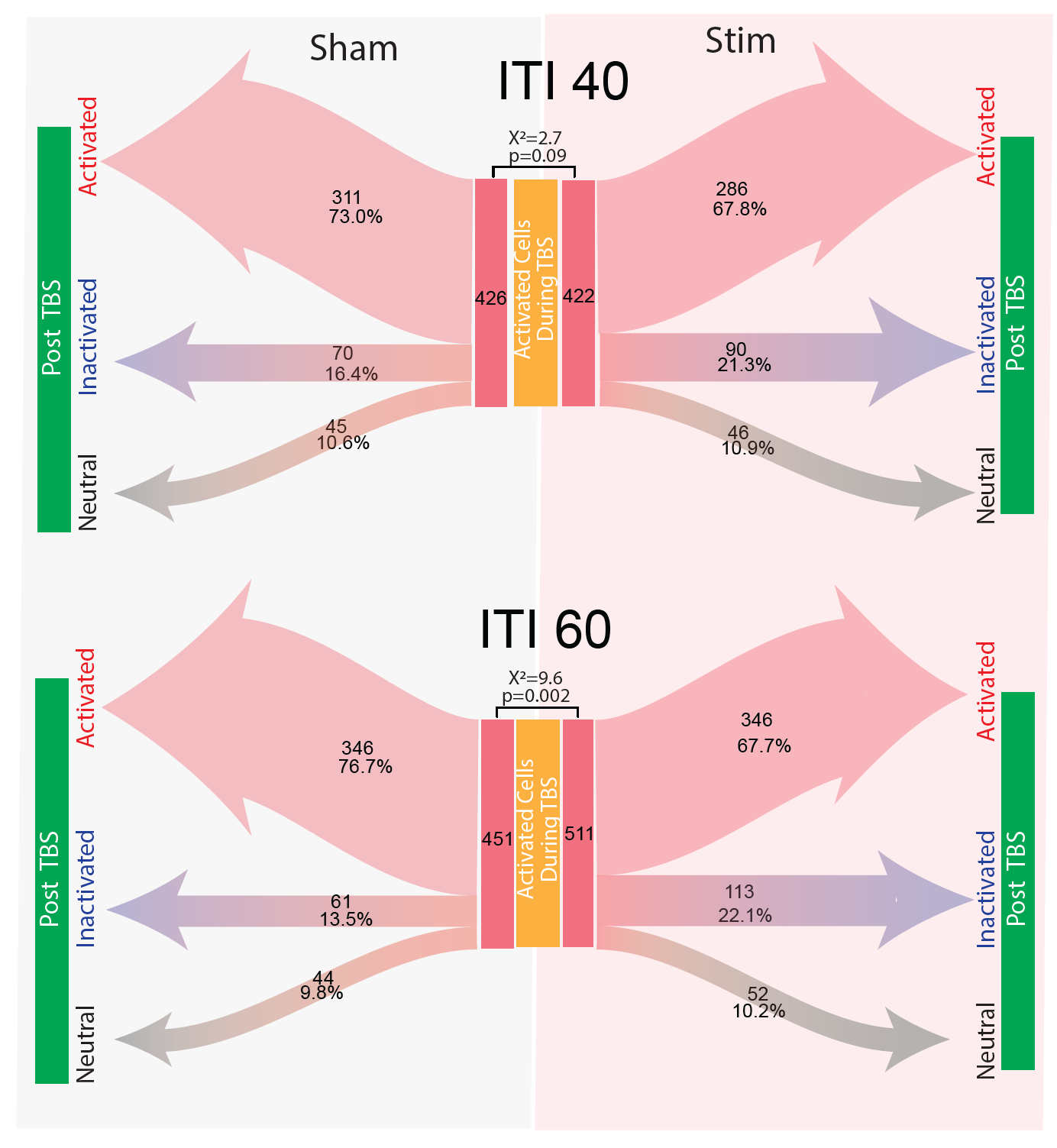

**Figure S4 . Stability of activated glutamatergic neurons is preserved at extended inter-train intervals.** Transition analysis of activated cells during stimulation was used to compare neurons that remained activated post-stimulation with neurons that transitioned to inactivated or neutral states. ITI 40 s did not significantly alter transition patterns relative to sham, indicating preservation of activated-cell stability after stimulation. At ITI 60 s, we found significance difference between activated post-stimulation with neurons that transitioned to inactivated or neutral states suggesting a partial reduction in response stability at the longest ITI.

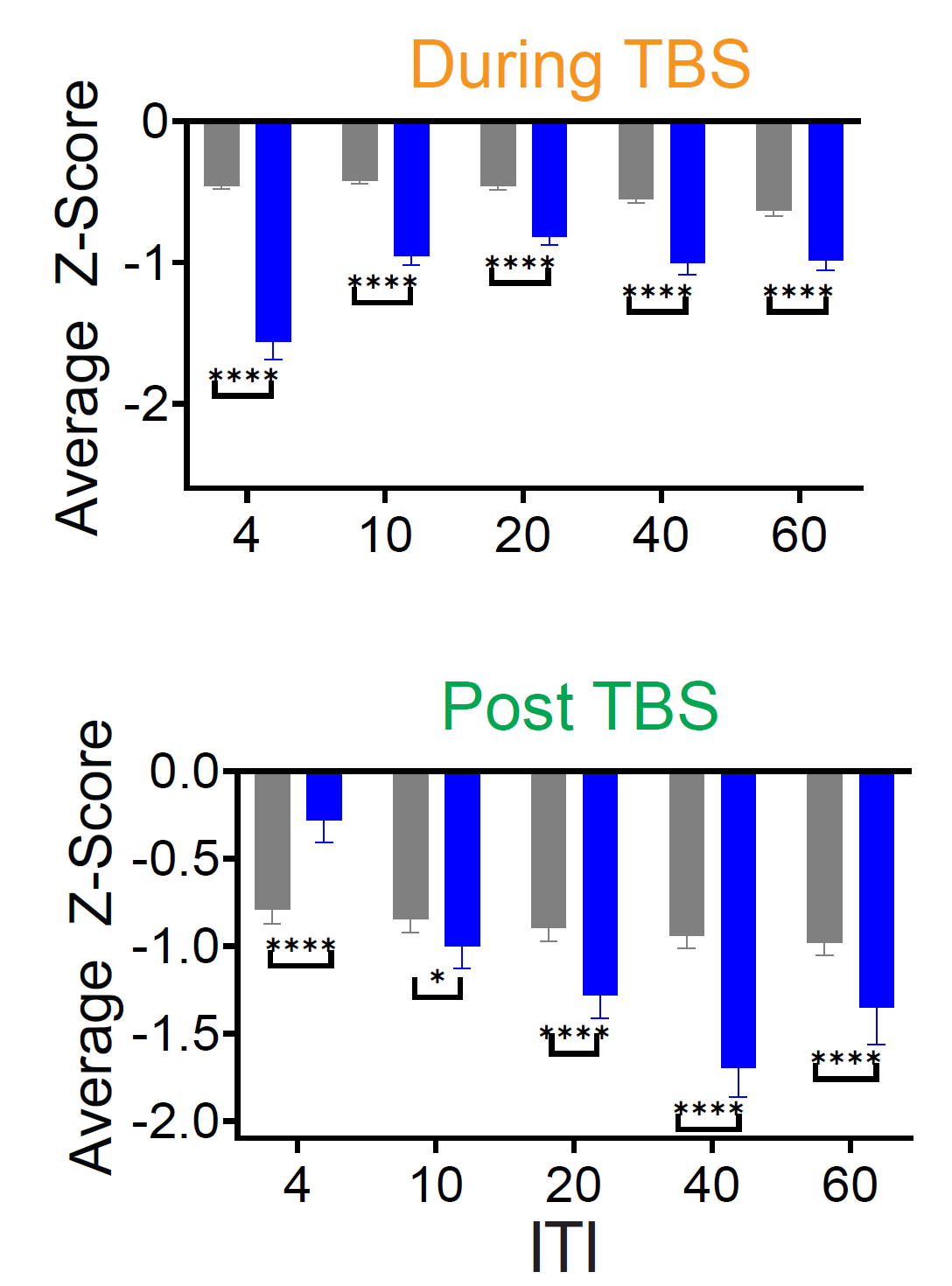

**Figure S5 . Inactivated GABAergic neurons.** Top panels show calcium activity during TBS, whereas bottom panels show post-stimulation activity. Average calcium activity of inactivated cells was compared between sham and stimulation conditions during and after TBS. During stimulation, inactivated cells exhibited significantly lower calcium activity than sham at all ITIs with the strongest suppression observed at ITI. Post-TBS, inactivated cells showed significantly higher activity than sham at ITI4, but lower activity than sham at 10-60s indicating that extended ITIs promoted sustained disinhibition after stimulation. Data were analyzed using two-way ANOVA followed by Sidak’s correction for multiple comparisons. Gray indicating sham and blue indicating stimulation condition. Values are presented as mean ± SEM

**Table S3. Comparison of Z-scored calcium activity in GABAergic cells during stimulation across inter-train intervals (ITIs).** Statistical analyses correspond to Figures 1D. For each ITI (4, 10, 20, 40, and 60 s), Z-scored calcium activity during stimulation was compared between sham and stimulation conditions using two-way ANOVA. Analyses were performed separately for all recorded GABAergic cells , activated neurons, and inactivated neurons. Reported values indicate the main effect of stimulation condition (sham vs. stimulation) within each ITI. Activated and inactivated neurons were classified based on significant deviations from baseline calcium activity during stimulation (p < 0.01).

|  | Cell Category in response to stimulation | | | |
| --- | --- | --- | --- | --- |
| ITI (Sec) |  | All | Active | Inactive |
|  | 4 | F=302.6  p<0.0001 | F=132.85  p<0.0001 | F=105.48  p<0.0001 |
|  | 10 | F=7.66  p=0.006 | F=8.93  p=0.003 | F=36.17  p<0.0001 |
|  | 20 | F=1.20  p=0.27 | F=2.38  p=0.12 | F=18.66  p<0.0001 |
|  | 40 | F=0.01  p=0.921 | F=0.23  p=0.625 | F=33.02  p<0.0001 |
|  | 60 | F=0.002  p=0.968 | F=0.202  p=0.65 | F=21.2  p<0.0001 |

**Table S4. Comparison of Z-scored calcium activity post-stimulation in GABAergic neurons across inter-train intervals (ITIs).** Statistical analyses correspond to Figures 3E. Post-stimulation calcium activity was quantified across eight consecutive 5-min bins spanning 40 min following TBS. For each ITI (4, 10, 20, 40, and 60 s), Z-scored calcium activity was compared between sham and stimulation conditions using two-way ANOVA. Analyses were performed separately for all recorded GABAergic neurons, activated neurons, and inactivated neurons. Reported statistics reflect the main effect of stimulation condition (sham vs. stimulation) within each ITI. Activated and inactivated neurons were classified based on significant deviations from baseline calcium activity during stimulation (p < 0.01).

|  | Cell Category in response to stimulation | | | |
| --- | --- | --- | --- | --- |
| ITI (Sec) |  | All | Active | Inactive |
|  | 4 | F=0.22  p=0.63 | F=37.64  p<0.0001 | F=30.45  p<0.0001 |
|  | 10 | F=7.2  p=0.007 | F=0.123  p=0.72 | F=4.11  p=0.043 |
|  | 20 | F=7.9  p=0.005 | F=0.93  p=0.33 | F=28.9  p<0.0001 |
|  | 40 | F=7.65  p=0.006 | F=0.35  p=0.55 | F=123.52  p<0.0001 |
|  | 60 | F=0.24  p=0.62 | F=0.073  p=0.78 | F=30.81  p<0.0001 |

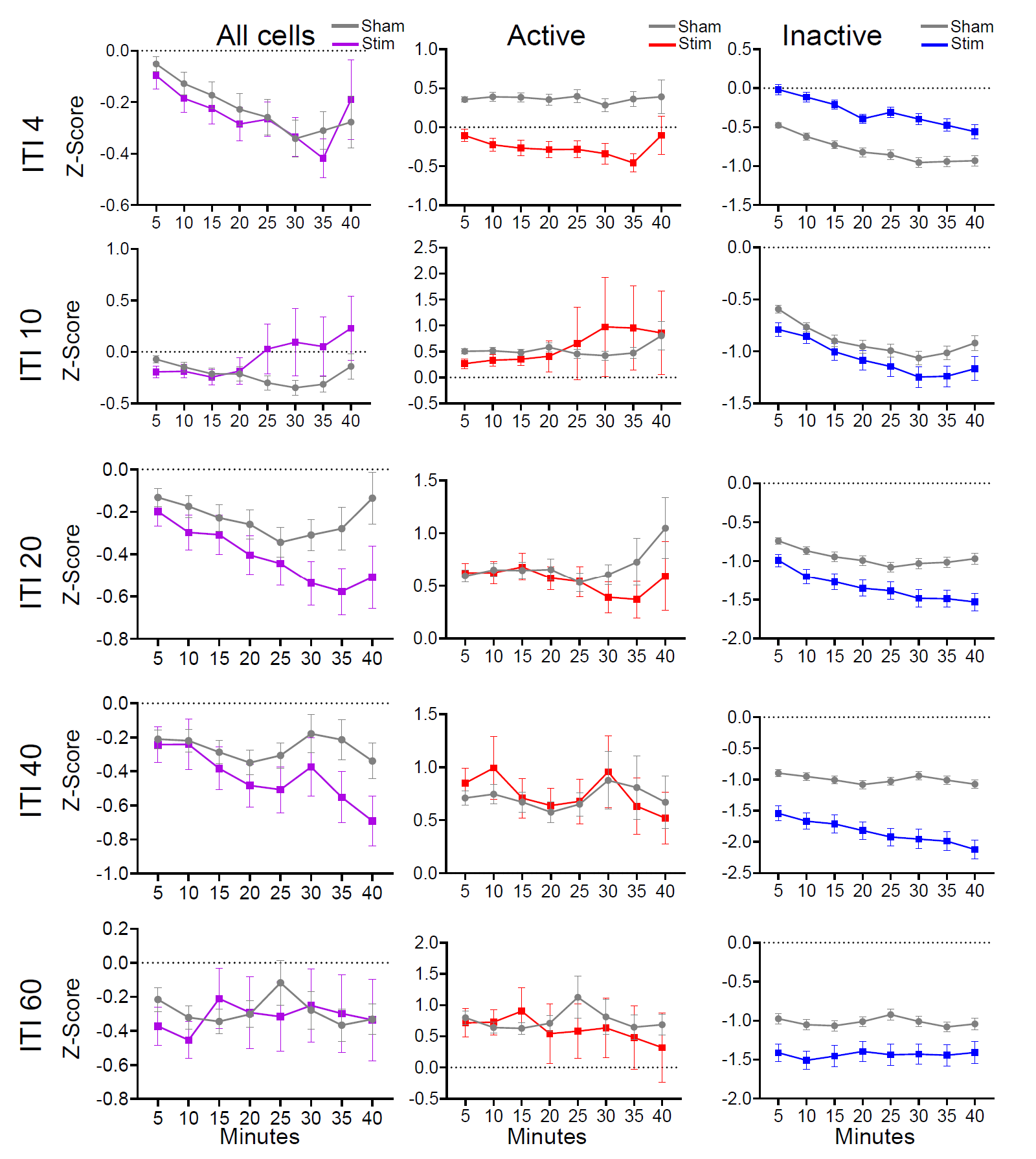
**Figure S6 . Comparison of Z-scored calcium activity post stimulation in GABAergic cells.** Post-TBS Z-scored activity 8 discrete of 5min for 40 min post time bins for all, active and inactive cells population. Each point indicates average of Z-score within 5 min time bin. shown mean ±SEM in corresponding fig 2 D. Z-scored calcium activity was calculated by normalizing activity across the session to this baseline.

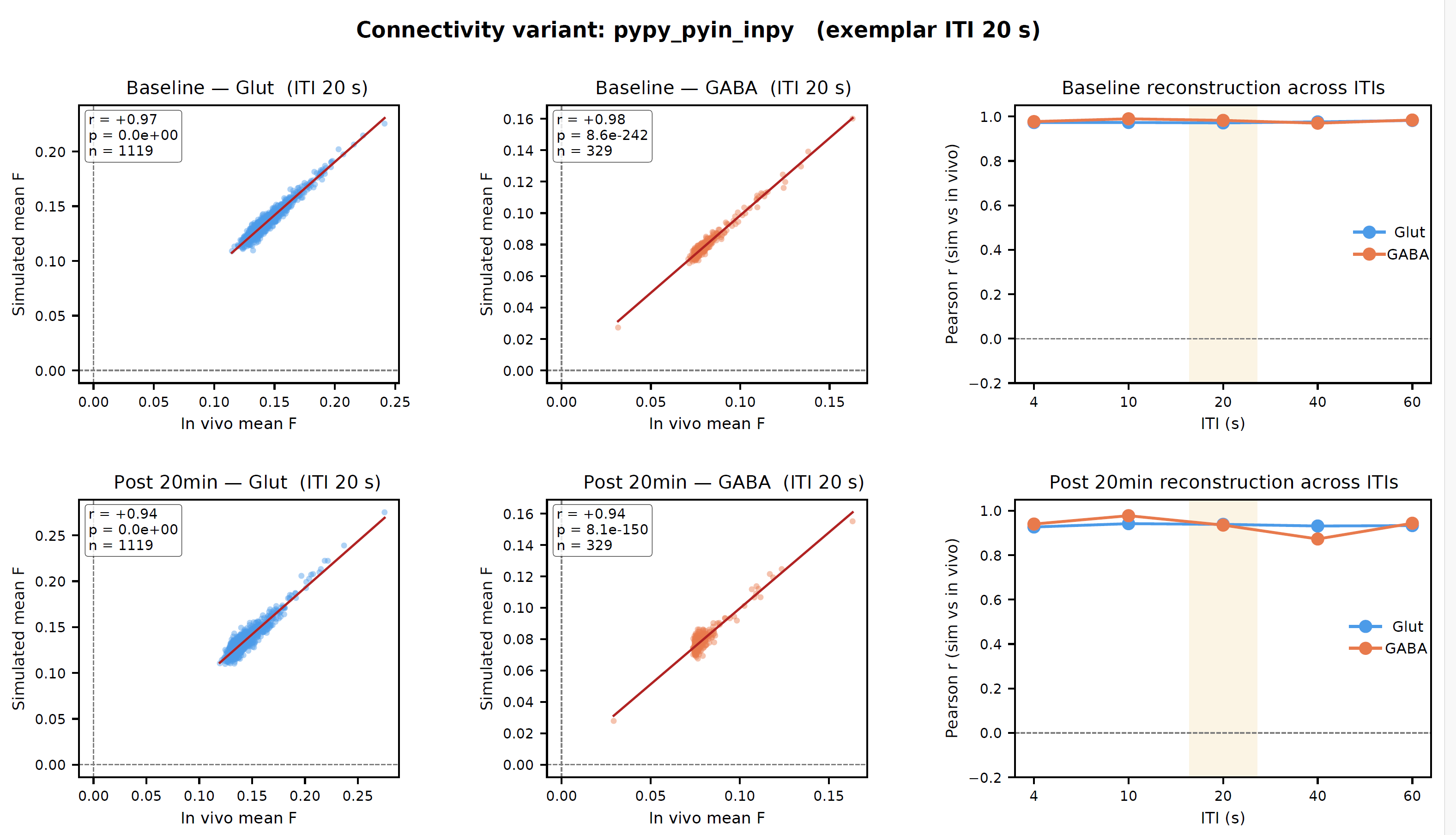

**Figure S7. Model reconstruction of in vivo single-neuron activity before and after TBS across ITIs.** Per-neuron average fluorescence in the simulated model is plotted against corresponding in vivo recordings for a representative inter-train interval (ITI; 20 s; left and middle columns) and summarized as Pearson’s r across all ITIs (4, 10, 20, 40, 60 s; right column). Each point represents a constrained neuron (glutamatergic, blue; GABAergic, orange), and the red line indicates the least-squares fit. The top row shows baseline activity prior to TBS, while the bottom row shows activity 20 minutes post-stimulation. In the summary panels, the shaded region denotes the representative ITI shown in the left panels. Reconstruction accuracy was high across all conditions (r > 0.9 for both cell types at all ITIs), indicating that the model robustly captures both baseline activity and TBS-induced reorganization of single-neuron dynamics.

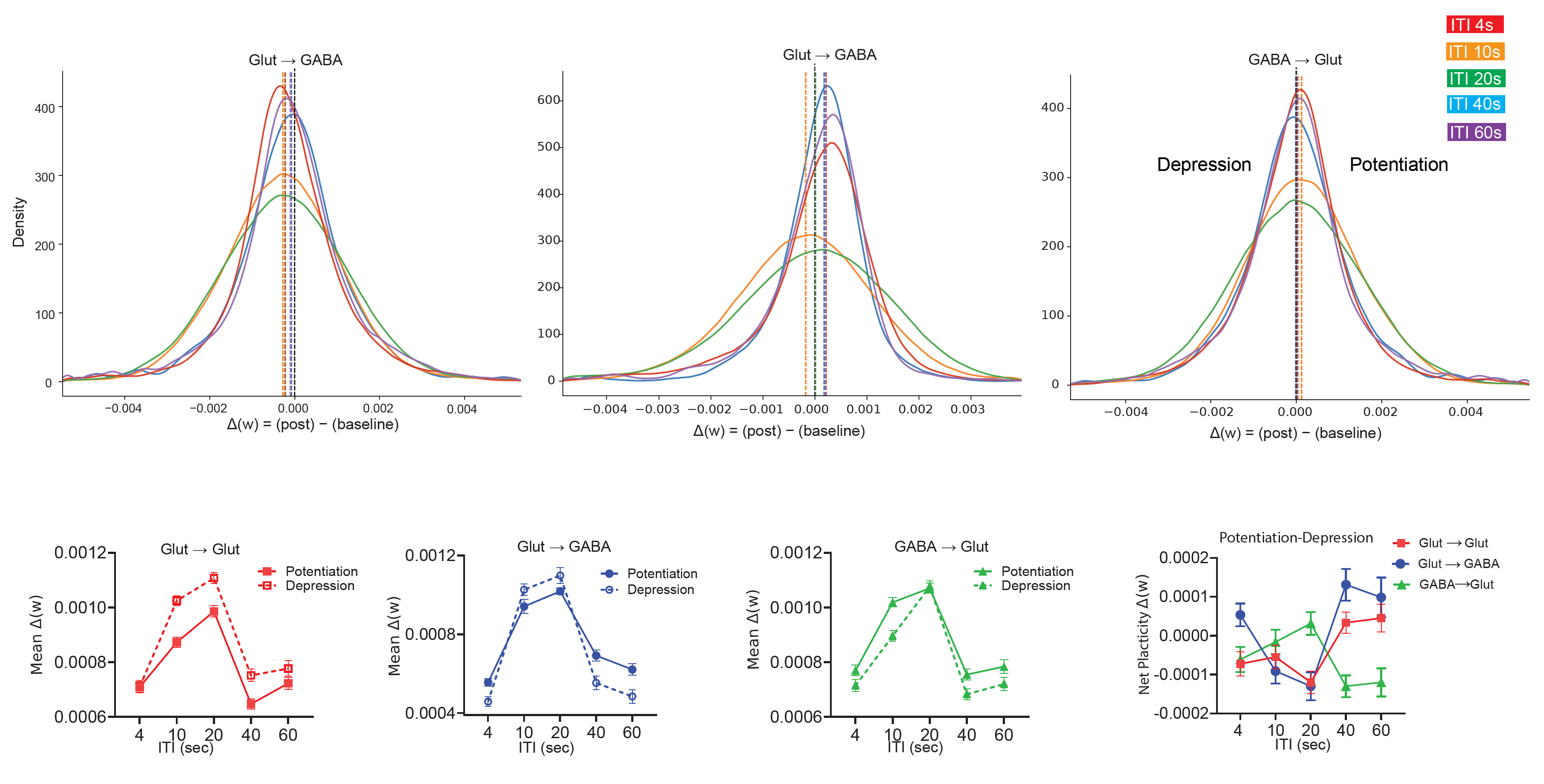

**Figure S8. Synaptic weigh in presence of GABA to glutamatergic connection across inter-train intervals (ITIs). (Top)** Distribution of model-derived synaptic weight changes (Δw; 20 min post-stimulation minus baseline) for connections between glutamatergic → glutamatergic; glutamatergic → GABAergic; and GABAergic → glutamatergic. **(Bottom)** Mean synaptic changes for glutamatergic → glutamatergic connections separated into potentiation and depression components. Two-way ANOVA revealed significant effects of synaptic state (potentiation vs. depression), ITI (p < 0.0001 for both), and their interaction (p= 0.0035), indicating ITI-dependent modulation of excitatory synaptic plasticity. Mean synaptic changes for glutamatergic-to-GABAergic connections showed similar significant effects for synaptic state (p=0.027), ITI and their interaction (p<0.0001 for both). Synaptic changes for GABAergic → glutamatergic connections showed ITI (p < 0.0001 for both), and their interaction (p= 0.036). Net synaptic plasticity (potentiation minus depression) revealed a dominant negative shift at ITI 20s (eTBS) in both glutamatergic → glutamatergic; glutamatergic → GABAergic connection and very negligible positive for GABAergic → glutamatergic connections.
